# Genome assembly and transcriptome analysis provide insights into the anti-schistosome mechanism of *Microtus fortis*

**DOI:** 10.1101/2020.09.03.282319

**Authors:** Hong Li, Zhen Wang, Shumei Chai, Xiong Bai, Guohui Ding, Junyi Li, Qingyu Xiao, Benpeng Miao, Weili Lin, Jie Feng, Cheng Gao, Yuanyuan Li, Bin Li, Wei Hu, Jiaojiao Lin, Zhiqiang Fu, Jianyuan Xie, Yixue Li

## Abstract

*Microtus fortis* (*M. fortis*) so far is the only mammal host that exhibits intrinsic resistance against *Schistosoma japonicum* infection. However, the underlying molecular mechanisms of this intrinsic resistance are not yet known. Here we performed the first *de novo* genome assembly of *M. fortis*, comprehensive gene annotation and evolution analysis. Furthermore, we compared the recovery rate of schistosome, pathological change and liver transcriptome between non-permissive host *M. fortis* and susceptible host mouse at different time points after Schistosome infection. We reveal that Immune response of *M. fortis* and mouse is different in time and type. *M. fortis* activates immune and inflammatory responses on the 10^th^ days post infection, involving in multiple pathways, such as leukocyte extravasation, antibody activation (especially IgG3), Fc-gamma receptor mediated phagocytosis, and interferon signaling cascade. The strong immune responses of *M. fortis* in early stages of infection play important roles in preventing the development of schistosome. On the contrary, intense immune response occurred in mouse in late stages of infection (28~42 days post infection), and cannot eliminate schistosome. Infected mouse suffers severe pathological injury and continuous decrease of important functions such as cell cycle and lipid metabolism. Our findings offer new insights to the intrinsic resistance mechanism of *M. fortis* against schistosome infection. The genome sequence also provides bases for future studies of other important traits in *M. fortis*.

## Background

Schistosomiasis is one of the most serious parasitic disease caused by blood flukes of the genus schistosoma. WHO estimates that at least 220.8 million people required preventive treatment for schistosomiasis in 2017 [1], thus schistosomiasis has a serious impact on health and economy [2]. Recent genome studies obtained the draft genomes of *S. japonicum*, *S. mansoni* and *S. haematobium*, providing insights into the complex mechanism of host-parasite interaction [3–6]. Schistosoma shares more orthologs with mammal hosts than those they share with ecdysozoans, which enables it to exploit the host’s metabolism and signal pathways to complete growth and development [3].

It was reported that *S. japonicum* could native infect 46 mammal hosts [7]. Schistosomes penetrate the skin of host, migrate through the heart and lung, and then develop in the liver. The development and survival of schistosomes were distinct among different hosts [7]. Around 40%~70% worms can complete their life cycle and cause severe pathological damage in susceptible hosts such as mouse, rabbit, cattle and goat. Only a small fraction of worms can survival in non-susceptible hosts such as rat and water buffalo (Supplementary Table 1). To our best knowledge, *M. fortis* is the only mammal in which schistosomes cannot get maturation [7]. The intrinsic resistance of *M. fortis* against schistosoma has been proved by multiple studies [8, 9], no matter *M. fortis* came from a schistosomiasis epidemic or a non-epidemic area [10]. Additionally, our previous studies demonstrate that *M. fortis* is also resistant to S. mansoni. Therefore, *M. fortis* is a valuable animal model to study the mechanism of host-schistosoma interaction.

*M. fortis* (reed vole) is a member of the Rodentia: Cricetidae order. It distributes in China, Korea, NE Mongolia and parts of Russia. Besides intrinsic resistance against schistosome, *M. Fortis* has potential to be animal models of human diseases, such as nonalcoholic fatty liver, diabetes and ovarian cancer [11, 12]. Due to the lack of genome sequence of *M. fortis*, many experiments had to use similar sequences from the close organism. For instance, mRNA microarray experiments used mouse microarray platform [13], and miRNA microarray studies were comprised of miRNAs in mouse, rat, and Chinese hamster [14]. Although these studies found some differential expressed genes, the results can not exactly present the molecular characteristics of *M. fortis*. A reference genome of *M. fortis* is in urgent need.

Previous studies have found several proteins that may be associated with the intrinsic resistance of *M. fortis* to *S. japonicum*. Heat shock protein 90α of *M. fortis* (*Mf*-HSP90α) caused 27.0% schistosomula death rate in vitro, and mice injected with Mf-HSP90α recombinant retrovirus reduced 40.8% worm burden [15]. Similar studies showed that mice injected with *Mf*-KPNA2 and *Mf*-albumin had 39.42% and 43.5% worm burden reduction, respectively [16, 17]. Another work reported that purified IgG3 antibody from laboratory-bred *M. fortis* and wild *M. fortis* could more effectively kill schistosomula than the IgG3 from Kunming mice [18]. Additionally, cytokines and chemokines levels in the sera of *M. fortis* are assessed to study the immune response changes. Expression of IL-4, IL-5 and IL-10 are increased from the second to the third week post-infection, indicating Th2 biased immune response is important for schistosoma clearance [19, 20]. Upregulation of IL-12 and interferon gamma (IFNγ) demonstrate the roles of Th1 immune response in *M. fortis* [19, 20]. However, most of these results have not been confirmed or been further investigated by other researchers. Investigation of one or several genes is insufficient to understand the complex interaction between schistosoma and *M. fortis*. In recent years, new technologies such as next generation sequencing are used to directly measure the molecular profiles of *M. fortis*. A study based on *de novo* transcriptome sequencing shows that innate and adaptive immune responses may play an important role in the intrinsic resistance against schistosome [21]. Another study used liquid chromatography-mass spectrometry to find some differential metabolites between infected *M. fortis* and C57BL/6 mice [19]. However, these large-scale studies cannot accurately locate the resistance genes due to lack of *M. fortis* genome. More studies are necessary to explore the mechanism of *M. fortis* intrinsic resistance.

Here we generate the draft genome of *M. fortis*, and annotate its genomic features comprehensively. The comparative transcriptome analysis reveals that non-permissive host *M. fortis* and susceptible host mouse are different in time and type of immune response. We propose key genes and pathways of immune response which serve as a basis for future experimental studies.

## Results

### Recovery rate of schistosomula from *M. fortis*

Although several studies have confirmed the intrinsic resistance of *M. fortis* to *S. japonicum* [7–9], the process of elimination of worms in their bodies is unclear. Therefore, we infected M. fortis and BALB/c mice with Schistosoma japonicum cercariae, and calculated the percentage of worms that were recovered by perfusion and culturing from different tissues of the infected animals. As shown in Figure 1A-B, the percentage of recovered worms (recovery rate) is significantly lower in infected *M. fortis* than that in infected mice. Total recovery rates of *M. fortis* are 15.04%, 7.05%, 9.47% and 0.77% on the 1st, 3rd, 7th and 14th days post infection (DPI) respectively, while those of BALB/c mice are 39.24%, 38.56%, 46.17%, 43.00%. The results indicate that schistosomula is extinct gradually in *M. fortis* in the early stages of infection (1-14 days). After infection for longer time (21^st^, 28^th^ and 42^nd^ DPI), schistosomula disappears in *M. fortis*, but there are still around 60% worms could be recovered from hepatic portal vein in BALB/c mice.

**Figure 1.**
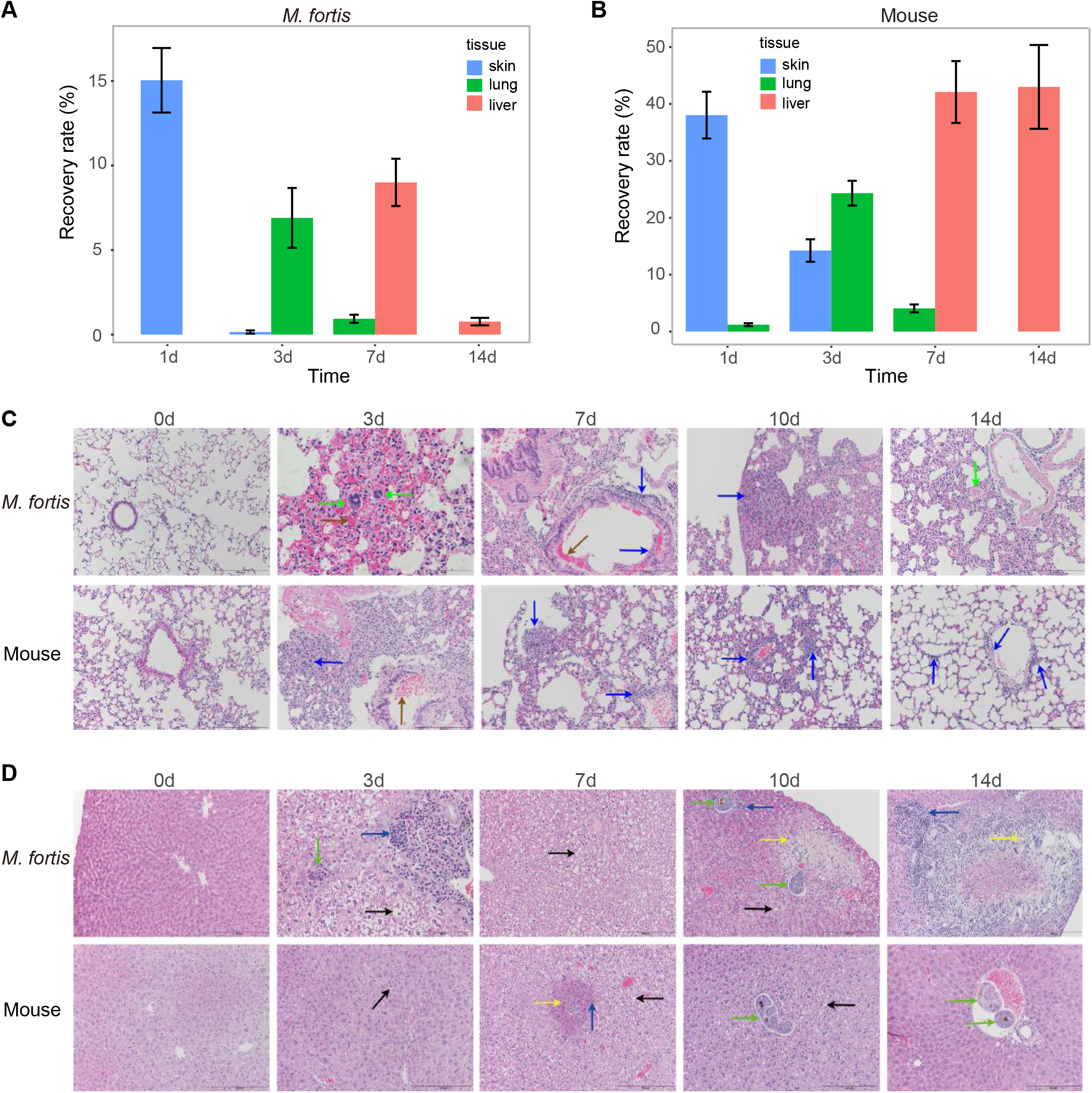
Phenotype of *M. fortis* and mouse after *S. japonicum* infection. A) Recovery rate of schistosoma from *S. japonicum*-infected *M. fortis*. B) Recovery rate of schistosoma from *S. japonicum*-infected mice. Schistosomula was obtained from skin, lung and liver of *M. fortis* and mice respectively. C) Histopathology of lung sections with HE staining. D) Histopathology of liver sections with HE staining. The up and down panels are the tissues prepared from *M. fortis* and mice before infection (0d) and on the 3^rd^, 7^th^, 10^th^ and 14^th^ days after *S.japonicum* infection (×200). Blue arrow stands for inflammatory cell infiltration, brown arrow stands for pulmonary hemorrhage, black arrow stands for liver cell vacuolation, green arrow stands for parasite, and yellow arrow stands for hepatocyte necro.

### Histopathological changes in the lungs and livers of *M. fortis* infected with *S. japonicum*

Some hemorrhagic spots were observed on the lung surface of *M. fortis* on the 3^rd^ and 7^th^ DPI, and the lung of BALB/c did not have similar damage (Figure 1C, Supplementary Figure1). Histopathological observation showed that the worms were found in the lung tissue of the *M. fortis* on the 3^rd^ and 7^th^ DPI, with large bleeding around them, and inflammatory cells infiltrating such as neutrophils and lymphocytes. There were similar pathological phenomena in the lungs of mice, but to a lesser extent.

Some white nodules which contained the remnants of schistosomula appeared on the liver of *M. fortis* beginning on the 7^th^ DPI, and most of them disappeared on the 14^th^ DPI (Figure 1D). The appearance of the liver of infected *M. fortis* returned to normal on the 21^st^ DPI. Histopathological observation showed that most of the remaining worms in nodules were in the small vessels, surrounded by inflammatory cell infiltration. There was no obvious pathological change in the liver of mice during the same period, although some schistosomula were *obser*ved in the liver slices. These histopathological changes were consistent with previous studies [22].

### Genome assembly and annotation

Genomic DNA of an 8-week-old female *M. fortis* from Dongting Lake, Hunan, China was subjected to shortgun sequencing (Table1, Supplementary Table 2). The sequencing depth was more than 110 X. Sequence reads were assembled by using ALLPATHS-LG into scaffolds. The final genome assembly was 2.2 Gb in length, which was about 92% of the estimated genome (Supplementary Figure 3, Supplementary Table 3). The contig N50 (the shortest length of sequence contributing more than half of assembled sequences) was 60.7 kb and the scaffold N50 was 10.1 Mb (Table1, Supplementary Table 4). The GC content was 42.4%, which was similar to that mammal genomes (Supplementary Figure 4, Supplementary Table 5). Assembly quality was assessed by CEGMA; a total of 238 core eukaryotic genes (96%) out of 248 were found in the assembly (Supplementary Table 6). Additionally, 116,254 transcriptional fragments (>200 bp) were identified by *de novo* RNA-Seq assembly and over 98% of them were covered by the assembled scaffolds (Supplementary Table 7).

The genome was analyzed for repeats and low complexity DNA sequences using RepeatMasker. The content of repetitive elements was 30.2%. It is lower than those reported for most mammalian genomes, but higher than that from another sequenced species *M. ochrogaster* (Prairie vole) in the Microtus genus (Supplementary Figure 5, Supplementary Table 8). The SINE, LINE and LTR repeats of *M. fortis* represent similar percentage (around 9%). The *M. fortis* and prairie vole genomes have about 35% fewer LINE-1 (387,111 and 386,270) than those the mouse and human genomes (617,477 and 579,553) have, suggesting that the LINE-1 repeat specifically decreased in the Microtus genus.

We identified 21,867 protein-coding genes by combining *de novo* prediction, homology-based prediction, and transcriptome-aided annotation (Supplementary Figure 6-8, Supplementary Table 9-11). Among protein-coding genes, 13,159 (60%) were identified from RNA-Seq data; 16,990 (78%) had homologs in other species; 21,128 (96.6%) genes could be annotated to protein families, KEGG pathways or Gene Ontology terms. We also identified 2,462 non-coding genes, including 670 miRNA, 515 tRNA, 167 rRNA, 147 lncRNA and 963 snRNA. The number and annotation of predicted genes are comparable to those of well-studied mammalian genomes.

### Evolution of gene families

In order to examine the genotypes underlying the adaptations of *M. fortis*, we constructed orthologous gene families, analyzed the expansion or contraction of gene families, and detected positively selected genes. We identified 23,575 single-copy orthologous families by using 18 mammalian genomes (Supplementary Table 12). There existed 19,070 orthologous families in four genomes (human, mouse, *M. fortis* and *M. ochrogaster*), 14,327 (75%) of which were shared by all, 448 were specific to the *M. fortis* genome (Figure 2A). We constructed a phylogenetic tree by analyzing the orthologous gene families (Figure 2B). *M. fortis* sat within rodents, and was the closest to *M. ochrogaster*. The divergence time between *M. fortis* and *M. ochrogaster* was 8.6 (95% CI: 4.2-15.5) million years ago. The ancestors of Microtus genus split from the ancestor of rats and mice approximately 46.3 (95% CI: 29.3-63.9) million years ago.

**Figure 2.**
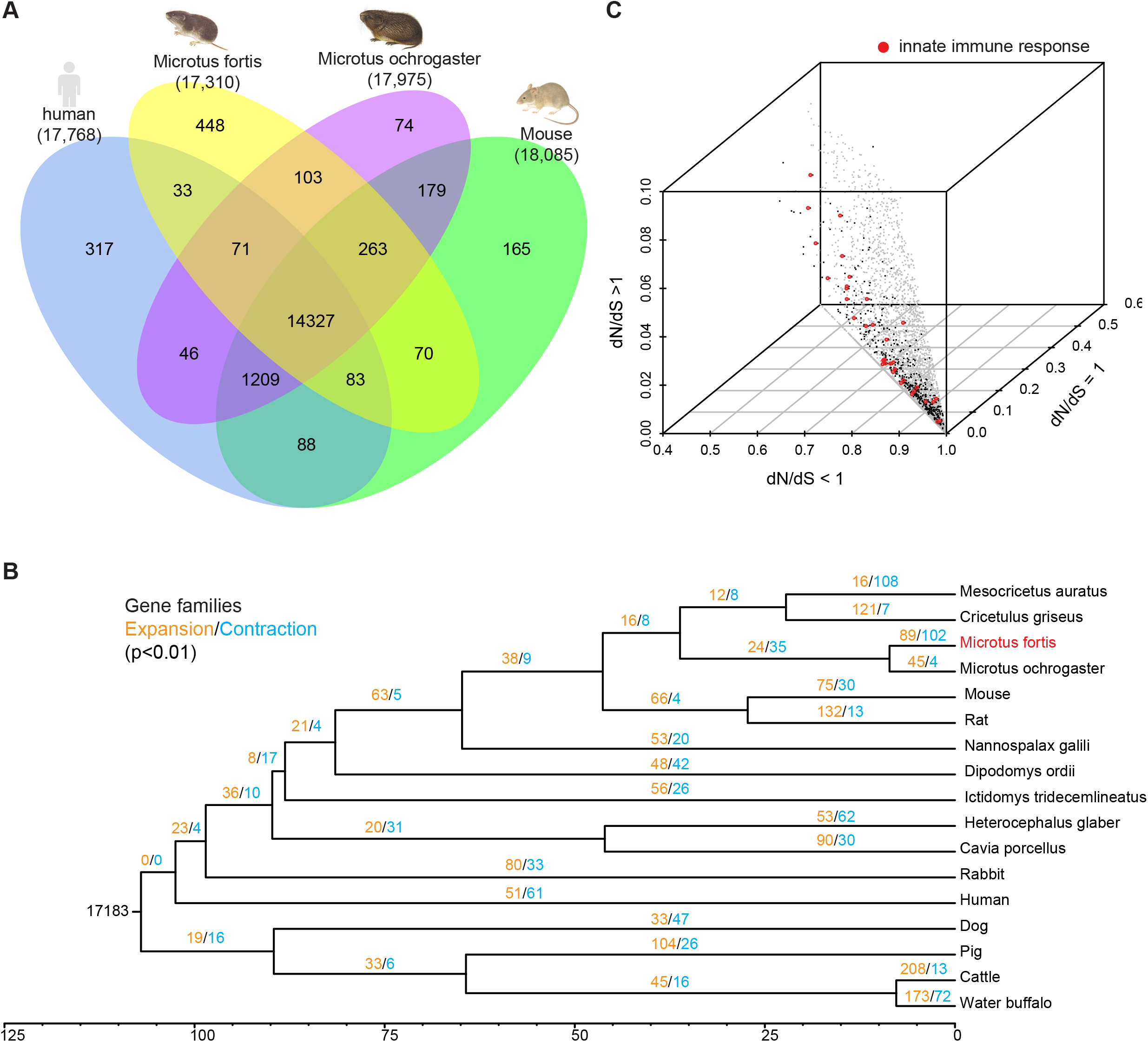
Analysis of *M. fortis* genome. A) A Venn diagram shows the unique and shared orthologous gene families in human, mouse, *M. fortis* and *M. ochrogaster*. B) Phylogeny tree and evolution of gene families. The numbers indicate the number of gene families that have expanded (orange) or contracted (blue) since the split from a common ancestor. C) dN/dS ratio of the positively selected genes. Three axes are the percentage of sites with dN/dS>1, dN/dS=1 and dN/dS<1, respectively. Genes involved in innate immune response (GO:0045087) are shown in red.

Compared to other mammals, *M. fortis* had 89 expanded and 102 contracted gene families. A large fraction of the contracted families involved olfactory receptors and taste receptors, which might be due to the limited food type of herbivore. We identified 532 positively selected genes (PSGs, Figure 2C) by employing the likelihood ratio test on the dN/dS ratio (ratio of the rate of nonsynonymous substitutions to the rate of synonymous substitutions). Functional enrichment analysis of 532 PSGs revealed rapidly evolving biological processes, such as regulation of cell shape (P=0.003) and innate immune response (P=0.004). A total of 33 innate immune genes were positively selected, which indicates the rapid evolution of immune system in *M. fortis* (Supplementary Table 13). For the previously reported genes (ALB[17], HSP90α [15], KPNA2[16]) that were possibly associated with *M. fortis* intrinsic resistance, we did not find any significant sites under positive selection.

We annotated the immunoglobulin (IG) and T cell receptor (TR) genes by aligning IMGT [23] reference sequences of human, mouse, rat and rabbit to the genome. For IG genes, we identified 122 (92) IGHVs, 146 (104) IGKVs and 33 (20) IGLVs (parentheses show sequences without stop codon) (Supplementary Table 14). For TR genes, we identified 92 (80) TRVAs/TRVDs, 34 (30) TRVBs and 6 (5) TRVDs. The TR loci were almost complete, with most V- and C-genes distributed in 4 scaffolds (Supplementary Table 15). Although the IG loci were more fragmented, our phylogenetic analysis of the IGV sequences suggested that they covered typical clan of other species such as human and mouse (Supplementary Figure 9). The IG and TR repertoire in the *M. fortis* genome provided valuable resources for screening specific immune molecules against Schistosoma.

Major histocompatibility complex (MHC) is a set of genes that are essential for immune system to recognize foreign molecule. We searched the Class I and II histocompatibility antigen domains across *M. fortis* genome. There were 27 genes with Class I histocompatibility antigen domains and 20 genes with Class II histocompatibility antigen domains (Supplementary Table 16). These genes were located on 11 scaffolds (>100kb). Seven scaffolds can be mapped to the MHC regions of mouse and human. (Supplementary Figure 10).

### Transcriptome characterization after *S. japonicum* infection

Since liver is the major organ where schistosome migrates, matures and dies, RNA-sequencing was performed to measure the liver transcriptome of *M. fortis* and mice before infection (0d) and at several time points related to key pathological changes (Supplementary Figure 11, Supplementary Table 17). We used “false discover rate (FDR) < 0.05 and the absolute value of fold change > 2” as threshold to select differentially expressed genes (DEGs) at different time points after *S. japonicum* infection. Compared to pre-infection (0d), 2,845 genes of *M. fortis* and 5,185 genes of mouse were differentially expressed at least one time point after infection (Supplementary Figure 12). *M. fortis*’s transcriptome changed mostly on the 14^th^ DPI, while mouse’s transcriptome changed mostly on the 28^th^ and 42^nd^ DPI. Generally, mouse had more DEGs than *M. fortis* had at the same time point, which is consistent with more severe pathological changes of mouse. For both species, a large number of DEGs overlapped at different time points after infection.

To investigate the dynamic expression pattern in time-series data, DEGs of *M. fortis* and mouse were divided into five and six subgroups based on hierarchical clustering (Figure 3AB, Supplementary Figure 14). Expression of genes in subgroup MF_C3 increased on the 7^th^ and 10^th^ DPI, then decreased on the 14^th^ DPI. Expression of genes in MM_C6 were stable until a significant increase on the 28^th^ and 42^nd^ DPI. Genes in these two clusters were especially interesting, since they are enriched in immune, inflammatory, defense response, cell adhesion and so on. Subgroup MM_C1 and MM_C3 represented the down-regulated genes on the 10^th^ DPI and 14^th^ DPI in mouse, which majorly participated in cell cycle, mitochondrion, lipid metabolism and some metabolic processes. MF_C1 was the largest subgroup of *M. fortis*, represent the up-regulated genes on the 14^th^ and 21^st^ DPI. However, there were no significant enriched functional terms in this subgroup.

**Figure 3.**
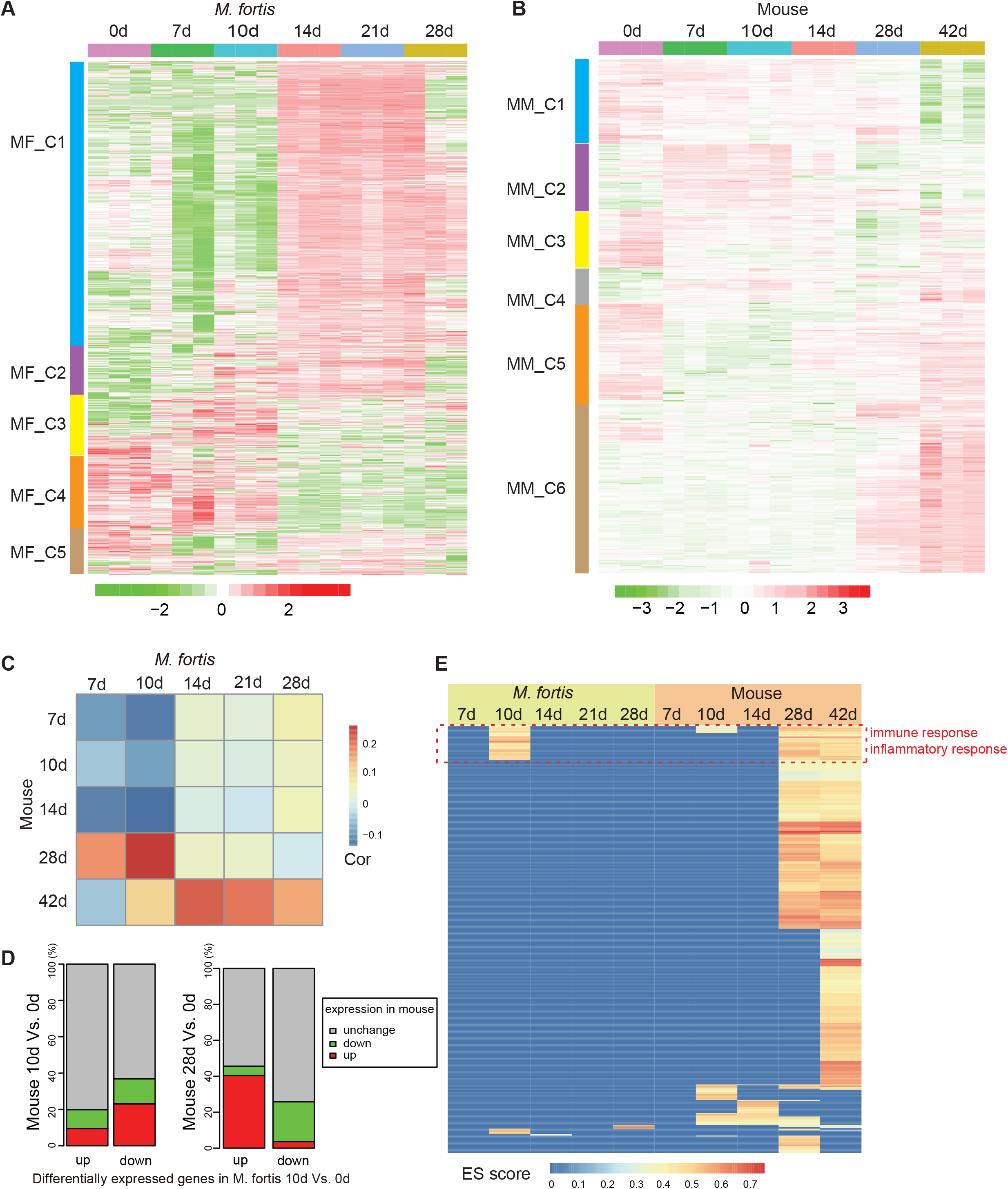
Comparative transcriptome analysis between *M. fortis* liver and mouse liver after *S. japonicum* infection. A) Time-series transcriptome of *M. fortis* liver before infection (0d) and on the 7^th^, 10^th^, 14^th^, 21^st^ and 28^th^ DPI. Rows were DEGs that were deferentially expressed between pre-infection and post-infection (FDR<0.05). Hierarchical clustering was used to classify the DEGs into clusters (The smallest cluster has more than 100 genes). B) Time-series transcriptome of mouse liver before infection (0d) and on the 7^th^, 10^th^, 14^th^, 28^th^ and 42^nd^ DPI. C) Correlation of gene expression changes. Expression fold changes of orthologous genes were calculated by comparing pre-infection (0d) and post-infection. Pearson correlation coefficients were calculated to measure the similarity of fold change between *M. fortis* and mouse. D) Comparison of the DEGs of *M. fortis* “0d Vs. 10d” with the DEGs of mouse “0d Vs. 10d”, and mouse “0d Vs. 28d”. E) Significantly enriched GO Biological Progress terms (FDR<0.05). The colors represent scores of Gene Set Enrichment Analysis. Positive (Negative) ES means gene expressions are up-regulated (down-regulated) after infection. GO terms in the red box were immune related biological process.

Comparative transcriptome analysis was used to explore the common and unique characterization of *M. fortis* and mice. Firstly, we compared DEGs between *M. fortis* and mouse at each paired time points. DEGs were obtained by comparing the expression profiles of post-infection with pre-infection (FDR<0.05). Correlation coefficients were calculated using the fold change of differentially expressed orthologs. The correlations between *M. fortis* and mouse were low at the same time point (Figure 3C), which indicated distinctive expression response to *S. japonicum* infection. Taken 10^th^ DPI as an example, 70% DEGs of *M. fortis* did not have significant expression change at mouse, and 92% DEGs at mouse were not differential at *M. fortis* (Figure 3D, Supplementary Table 18). Only 47 (100) genes were simultaneously up- (down-) regulated in both species on the 10^th^ DPI. The highest correlation occurred on the 10^th^ DPI of *M. fortis* and the 28^th^ DPI of mouse. 40% of the up-regulated genes on the 10^th^ DPI in *M. fortis* also increased significantly on the 28^th^ DPI in mouse (Figure 3D).

Secondly, we compared the annotated functions of DEGs in *M. fortis* and mouse by Gene Set Enrichment Analysis. Figure 3E Illustrated the significantly differential biological progress at different time points post-infection compare to pre-infection (Supplementary Figure 13). The enrichment of DEGs in *M. fortis* was majorly on the 10^th^ DPI, while DEGs in mouse enriched in more biological process terms on the 28^th^ DPI and 42^th^ DPI. Most of the significantly changed functions on the 10^th^ DPI of *M. fortis* were also enriched on the 28^th^ and 42^th^ DPI of mouse. The shared terms were majorly immune-related processes, such as immune system response, response to external stimulus and inflammatory response (Figure 3E, Supplementary Figure 13). These results demonstrated that intense immune responses of M. fortis occurred on the 10^th^ DPI. Although mouse had mild immune response after S. japonicum infection [24], more intense immune responses occurred at the 28^th^ and 42^th^ DPI in mouse.

### Distinct immunity mechanism of *M. fortis* and mouse

The above results showed that the 10^th^ day after infection is especially important to understand the differential immune response of *M. fortis* and mouse against *S. japonicum* infection. Therefore, we analyzed the annotated functions of 395 specially up-regulated genes on the 10^th^ DPI of *M. fortis*. Most of the enriched pathways were related with innate immunity. The top 3 pathways were leukocyte extravasation signaling, integrin signaling, and Fc-gamma receptor mediated phagocytosis in macrophages and monocytes (Figure 4A). ICAM1 (CD54) and VCAM1 (CD106) are important cell adhesion molecules in leukocyte extravasation signaling (Figure 4B). Their up-regulation may promote leukocytes migrate from the blood vessel to liver to eliminating schistosomulum. Expression of FCGR1 (CD64) increased on the 10^th^ DPI in *M. fortis* (Figure 4C). FCGR1 is the high-affinity receptor for IgG. It involved in phagocytosis and regulation of cytokine production. We further analyzed the expression pattern of immunoglobulin isotypes. IgG1, IgG3, IgA and IgM were partially up-regulated in *M. fortis* on the 7^th^ and 10^th^ DPI, significantly up-regulated on the 14^th^ and 21^th^ DPI (Figure 4D). On the contrary, expression of IgG1, IgA and IgM increased significantly in mice on the 28^th^ and 42^th^ DPI (Figure 4D). IgG3 is the most interesting antibody, since it was significantly up-regulated in infected *M. fortis*, but it almost did not increase in infected mouse except several outlier data points (Figure 4E). This result supports a previous study that IgG3 antibody purified from *M. fortis* could more effectively killed schistosomula than that of mouse [18].

**Figure 4.**
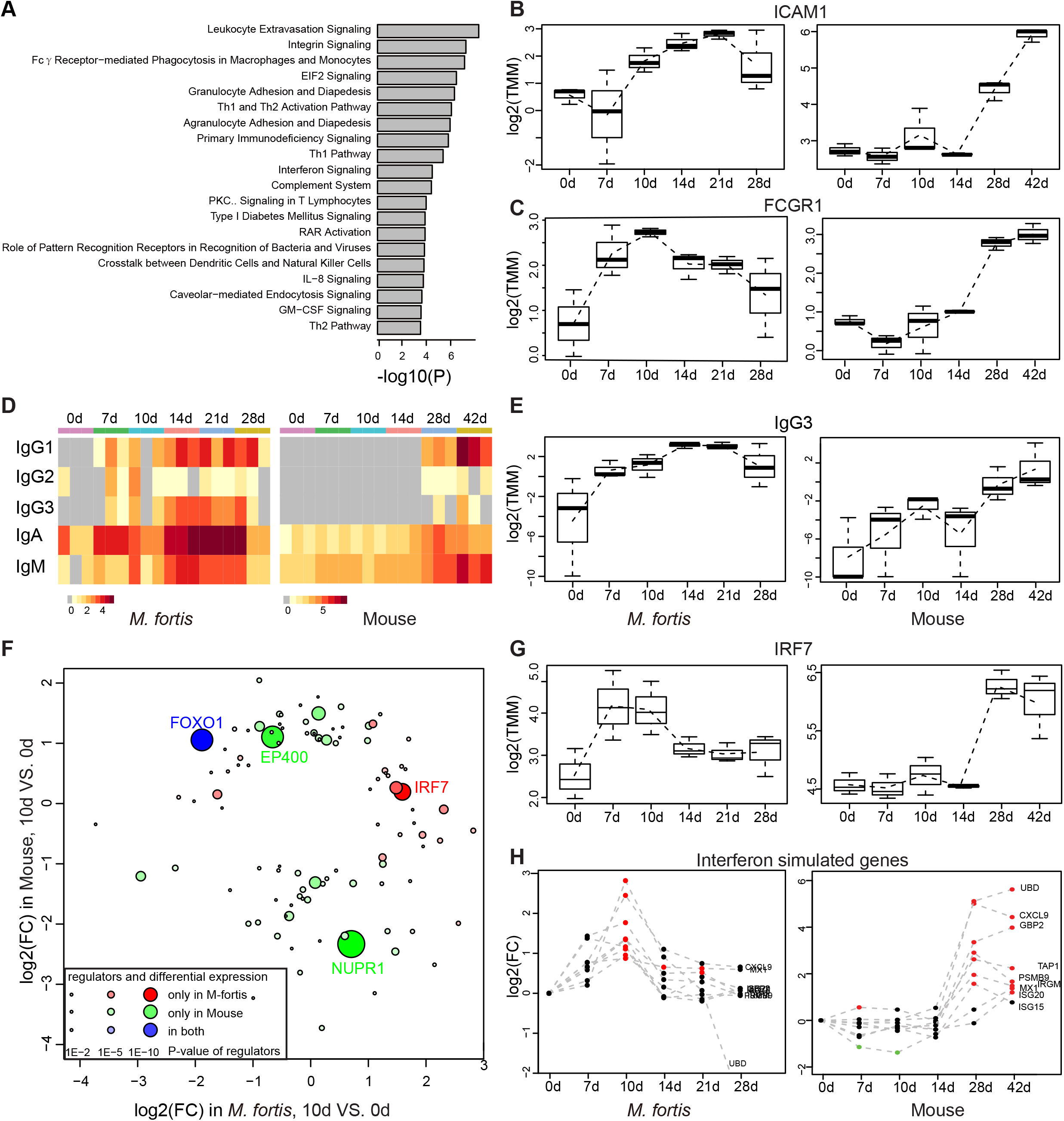
Mechanism of immune response in *M. fortis* against schistosoma infection. A) Enriched pathways of 395 genes that were up-regulated in *M. fortis* and unchanged in mouse. B), C) Expression profiles of ICAM1 and FCGR1. The expression value is given by log2(TMM), which were obtained from RNA sequencing. D) Expression change of immunoglobulin isotypes. E) Expression profiles of IgG3. F) Predicted regulators of *M. fortis* and mice on the 10^th^ days post-infection. Upstream regulators of the DEGs were identified by Ingenuity Upstream Regulator Analysis. Here we only showed regulators whose expression significantly changed in *M. fortis* or mouse, and the direction of expression change were not consistent between two species. Red and green circles indicate regulators that are specific for *M. fortis* and mouse, respectively. Blue circles represent the common regulators whose expression changes are opposite in *M. fortis* and mouse. G) Expression profiles of IRF7. H) Expression change of interferon simulated genes. Red and green points indicate significant up-regulation or down-regulation, respectively.

Furthermore, we used Ingenuity Pathways Analysis (IPA) to identify the potential upstream regulators on the 10^th^ DPI, which may explain the expression change of other genes. To remove false positive regulators, we only kept regulators which were differentially expressed at the corresponding time points. We supposed that the species-specific regulators or regulators with opposite expression change in two species were more important. In the end, we obtained 34 *M. fortis* specific regulators and 65 mouse specific regulators (Figure 4F). IRF7 was the top significant regulator in *M. fortis* on the 10^th^ DPI (P=9.03E-11). Further analysis of regulators on the 7^th^ DPI also revealed that IRF7 ranked the first in *M. fortis* (P=3.27E-8). IRF7 encodes interferon regulatory factor 7, is the major transcription factor that regulate type I interferon [25]. The expression of IRF7 significantly increased in *M. fortis* on the 7^th^ and 10^th^ DPI, and then decreased (Figure 4G). However, IRF7’s expression unchanged in mice in the first 10 days and then increased since the 28^th^ days post-infection (Figure 4G). To confirm the functional effect of IRF7 up-regulation, we analyzed interferon stimulated genes (ISG) whose expression are induced by interferon [26, 27]. As expected, expression pattern of ISGs were consistent with the expression pattern of IRF7 in both *M. fortis* and mouse (Figure 4H). We also observed the activation of JAK-STAT signaling pathway and up-regulated of some cytokines. Therefore, IRF7 was an important factor that activated the interferon signaling in *M. fortis* on the 7^th^ or 10^th^ DPI.

We used the same methods to analyze the mouse transcriptome on the 28^th^ DPI. Th1 and Th2 activation pathway was rank first in the IPA pathway analysis. The top two regulators were IFNG and TNF. Both genes are critical inflammatory cytokines of Th1 cells. Their expression values were significantly upregulated in mouse on the 28^th^ DPI, suggesting the activation of Th1 immune response (Figure 5A). This observation is consistent with the previous report that Th1 is the dominant immunity in the first 3–5 weeks in the mouse model of schistosome infection [24]. Although the immune response in mouse is strong since the 28^th^ days post-infection, it cannot eliminate schistosoma. Earlier intense immune response of *M. fortis* prevents the development of schistosomulum.

**Figure 5.**
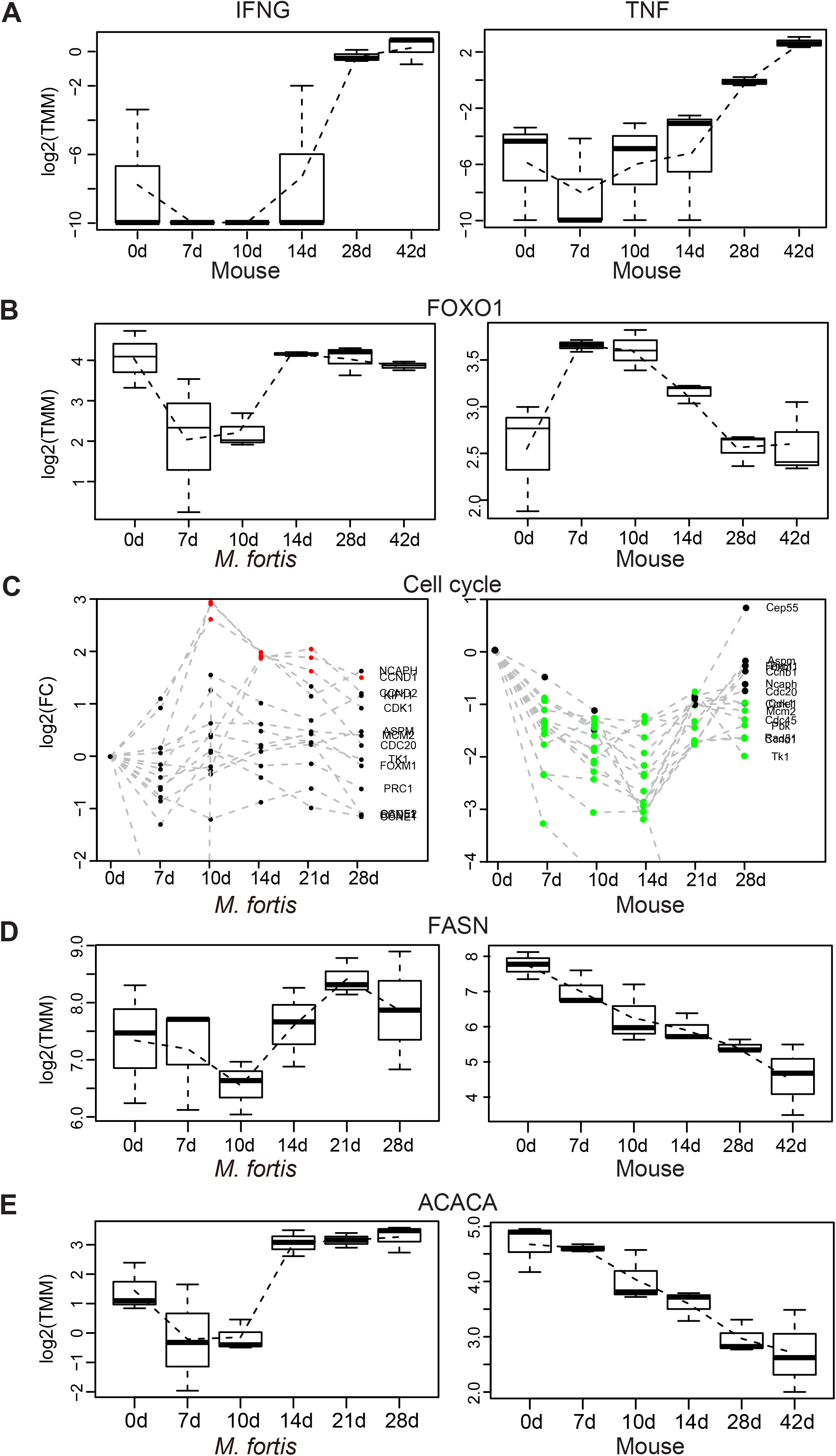
Transcriptome characteristics of mouse infected by schistosome infection. A) Expression profiles of IFNG and INF in mouse. B) Expression profiles of FOXO1 in *M. fortis* and mouse. C) Expression change of key genes involved in cell cycle. D), E) Expression profiles of FASN and ACACA.

### *S. japonicum* induced dysfunction in mouse

Since schistosoma infection induces severe pathological changes, we are also interested in the dysfunction of mouse. Transcriptome analysis revealed significantly down regulation of cell cycle and lipid metabolism on the 10^th^ days post infection. The regulator analysis showed that FOXO1 is the most significant gene that predicted to be common regulators of *M. fortis* and mouse (Figure 4F); Its expression decreased in *M. fortis* on the 7^th^ and 10^th^ DPI, while increased in mouse on the 7^th^ and 10^th^ DPI (Figure 5B). FOXO1 belongs to the forkhead family of transcription factors. It plays important roles in glucose and lipid metabolism, cell cycle arrest, and inflammation [28, 29]. Previous report showed that FOXO1 inhibited cyclin dependent kinases [30], therefore the over-presentation of down-regulated cell cycle genes in mouse may be the result of FOXO1’s activation (Figure 5C).

Schistosoma cannot *de novo* synthesize fatty acids [31]. We noted that fatty acid biosynthesis in mouse significantly decreased on 10^th^ day post schistosoma infection. Two key genes (FASN and ACACA) exhibited continuously decreasing expression pattern in schistosoma-infected mouse (Figure 5D,E), which is consistent with a previous report that major fatty acids and tricarboxylic acid cycle intermediates were significantly reduced in mice [32]. However, the expression of FASN and ACACA in *M. fortis* decreased on the and 10^th^ DPI and then returned to the original levels. Distinct expression pattern of fatty acid synthesis indicated different host-parasite interaction.

## Methods

### Population establishment

*M. fortis* (*Microtus Fortis* calamorum Thomas) were captured in Dongting Lake region, Hunan Province by the Institute of Subtropical Agriculture (ISA), the Chinese Academy of Sciences in 1994 and were bred in laboratory. *M. fortis* were introduced to Shanghai SIPPR-BK

Laboratory Animal Co., Ltd from ISA in 1998. Outbreed *M. fortis* colony was established [33, 34] and the voles were purified by biological purification [35]. The study protocol was approved by the Animal Care and Use Committee of the Shanghai Laboratory Animal Research Center.

### Genome sequencing and assembly

An 8-week-old female *M. fortis* was acquired after four generation inbreeding, and was subjected to genome sequencing. Genomic DNA was isolated from muscle, liver, lung and blood using Qiagen DNeasy kit according to the manufacturer’s instructions. Seven different paired-end libraries were constructed with 250 bp, 500 bp, 2 kb, 5 kb, 10 kb and 20 kb) insert sizes. The sequencing was done using Illumina HiSeq X Ten system in 2*150bp paired-end mode. The raw data were filtered to trim reads with adaptor sequences and remove low-quality reads. The remained reads were used to complete the genome assembly using ALLPATHS-LG. In order to assess the assembly quality, we used CEGMA (core eukaryotic gene mapping method) to identify the core genes in the *Microtus fortis* genome assembly.

### Genome annotation

RepeatMasker (v4.0.6) [36] was used to screen and annotate repetitive elements. Results from *de novo* repeat discovery by RepeatModeler (v1.0.8) [37] and homologous search against rodentia repeats in Repbase (v16.10) [38] were combined and masked. The repeat information of other genomes for comparison were fetched from RepeatMasker datasets online (http://repeatmasker.org/genomicDatasets/RMGenomicDatasets.html). Gene models were predicted by three approaches: 1) *de novo* prediction was performed on the repeat masked genome by five programs: AUGUSTUS (v3.0.1), GENEID (v1.4.4), GeneMark_ES (v2.3e), GlimmerHMM (v3.0.2) and SNAP (v2013-11-29). 2) homology-based prediction by projecting protein sequences of other mammals from RefSeq to the new genome. Rough search was performed by genBlastA (v1.0.1) [39], with protein coverage greater than 30%. Precise projection aware of gene structure was then performed by GeneWise (v2.4.1) [40] for the targeted DNA sequences. 3) Transcriptome-aided annotation was done by mapping RNA-seq reads back to the assembled genome using Tophat and Cufflinks. In the end, genes obtained from all the three approached were merged by the EVidenceModeler algorithm using a weight combination (*de novo* predictions = 0.3, GeneWise = 5, transcriptome = 10). Those EVM predictions supported by only one *de novo* program were removed, and predictions with a coding score below 1024 or a coding/noncoding score ratio below 2 are eliminated (supplementary Figure6). Pseudogenes were predicted by PseudoPipe which aligned the human proteins to *M. fortis*’s genome and reported a set of good-quality pseudogene sequences based on a combination of criteria [41].

Four types of noncoding RNAs (microRNAs, transfer RNAs, ribosomal RNAs and small nuclear RNAs) were annotated using tRNAscan-SE (v1.3.1) and Rfam database (v11.0). InterProScan (v5.19) was used to screen proteins against multiple protein signature databases, such as Pfam and Prosite. The KASS server was used to assign genes to KEGG ortholog and pathway. Gene ontology (GO) terms of human genes and mouse genes were assigned to their orthologs in *M. fortis*.

### Gene family construction

Gene families were constructed following the TreeFam pipeline [42], as described in Li et al.[43]. Protein sequences of other species were downloaded from RefSeq, and the longest isoform of each gene was preserved. Pairwise all-to-all blast were performed with e-value of 1e-10. Local alignments were joined by solar, and the total alignment length should cover at least 1/3 on both proteins. A h-score was calculated for each protein pair (p1, p2) based on the blast score: h-score = score (p1, p2)/max(score(p1, p1), score(p2, p2)). Homologous proteins were then clustered by hcluster_sg with minimum edge weight of 5, minimum edge density of 1/3 and opossum as an outgroup. For each cluster, multiple alignment on protein sequences was done by clustalo (v1.2.0) [44], which was then translated back to CDS alignment by treebest backtrans. Guided by the common tree from NCBI Taxonomy, the phylogenetic tree for each cluster was constructed by treebest best. Orthologs were inferred from the cluster by treebest nj. Solar, hcluster_sg and treebest were obtained from https://sourceforge.net/p/treesoft/code/HEAD/tree/branches/lh3/. Four-fold degeneration sites were extracted from the CDS alignment of single-copy orthologs, which were used to reconstruct the phylogenetic tree of species by MEGA (v7.0.18) [45]. The species tree was calibrated by MCMCtree in PAML (v4.9) [46], taking the divergence time (2.5% lower and upper bounds) of mouse-rat (11-47 Mya), mouse-human (67-124 Mya) and mouse-dog (65-150 Mya) from TimeTree [47]. Evolution of gene family size was inferred by CAFÉ (v3.1) [48] based on the homologous clusters. For families with significant size variations (family-wide p-value < 0.01), the branches with significant expansion and contraction were selected (Viterbi p-value < 0.01).

### Positively selected genes

Based on the CDS alignment of single-copy orthologs, positively selected genes in *M. fortis* were identified by codeml in PAML (v4.9) [46]. Poorly aligned regions were filtered by Gblocks (0.91b) [49]. Taking *M. fortis* as foreground and six schistosome-susceptible hosts (human, dog, cattle, mouse, rabbit, golden hamster) as background, the branch-site model (model = 2, NSsite = 2) with dN/dS ≤ 1 (fix_omega = 1, omega = 1) and dN/dS > 1 (fix_omega = 0) were adopted, respectively. The genes with significant dN/dS > 1 were identified by the likelihood ratio test (P < 0.05, chi-square test), and the positively selected sites were identified by the Bayes Empirical Bayes analysis.

### Immunoglobulins and T cell receptors

For the immunoglobulins (IG) and T cell receptors (TR), IMGT [23] reference sequences of germline V-, D-, J-, C-genes from the human, mouse, rat and rabbit were downloaded (release 201839-3). A rough alignment of amino acid sequences of V- and C-genes to the genome was performed with tblastn -e 1e-5. Redundant hits were merged, and V-segments with length short than 200 bp were filtered. The targeted regions were extracted for precise alignment with Exonerate (v2.2.0) [50] --model protein2genome --percent 50. For the scaffolds that contain V- or C-genes, the D- and J-genes were further mapped with tblastn. The V-genes were also annotated with the best hit to human or mice by IgBlast (v1.10.0) [51]. Multiple alignment of V-genes was performed by ClustalW, and the neighbor-joining tree was constructed by MEGA7 with default parameters.

### Infection Experiment

*S. japonicum* cercariae were harvested from positive *Oncomelania hupensis* snails maintained by Shanghai Veterinary Research Institute, Chinese Academy of Agricultural Sciences (Shanghai, China). BALB/c mice (male, 6 weeks old) and *Microtus fortis* (male, 6 weeks old) were provided by Shanghai SIPPR-BK Laboratory Animal Co., Ltd. Animal experiments were performed according to the protocols approved by the Animal Care and Use Committee of the Shanghai Veterinary Research Institute, Chinese Academy of Agricultural Sciences. 20 *M. fortis* voles and 20 BALB/c mice were randomly divided into 4 groups of 5 each and infected with 500±5 cercariae through the shaved abdominal skin. Schistosomula were collected from skin (2cm×2cm at infection site), lung or liver of infected animals by perfusion method and tissue culture on the 1^st^, 3^rd^, 7^th^ and 14^th^ days post-infection, respectively. The worm recovery rate was calculated as follows: percent of recovery = number of schistosomula /number of cercariae challenged ×100%.

### Histopathological assessment

15 *M. fortis* voles and 15 BALB/c mice were subdivided into five groups of 3 each, and animals in the four groups were percutaneously infected with 200±2 cercariae. Animals in each group were sacrificed either before infection or on the 3^rd^, 7^th^, 10^th^ and 14^th^ days post-infection. Lung and liver tissues were subjected to histopathological section analysis. Preparation of paraffin sections and histological assessment were executed by Shanghai SIPPR-BK Laboratory Animal Co., Ltd. Sections were stained with hematoxylin and eosin and observed using a light microscope (Nikon, Japan).

### RNA sequencing

*M. fortis* liver tissues were collected before infection (0d) and at 7^th^, 10^th^, 14^th^, 21^th^ and 28^th^ DPI. Mice liver tissues were collected before infection and at 7^th^, 10^th^, 14^th^, 28^th^ and 42^th^ DPI. There were three biological repeats at each time point. RNA was exacted and preserved in RNAlater^®^ (Ambion) at –80°C for RNA sequencing. Additionally, RNA was isolated from multiple tissues (heart, liver, lung, and kidney) from one *M. fortis*, and the mixed RNA was sequenced to identify more transcripts. The sequencing Libraries were constructed using the standard protocols, and were sequenced using 2×150bp paired-end strategy with Illumina HiSeq X Ten platform. Adapter sequences and low-quality bases were removed or trimmed by NGS QC Toolkit (v2.3.3). To assess the quality of genome assembly, RNA-seq data from all *M. fortis* samples were *de novo* assembled into transcriptional fragments by Trinity (v2.1.1). We then assessed the coverage of the transcripts in the genome assembly by mapping the assembled transcriptional fragments to the genome assembly using BLAT.

### Transcriptome analysis

RNA-Seq data of *M. fortis* and mouse were respectively mapped to the assembled draft genome and mouse genome (mm10) by RSEM algorithm (v1.2.21). Gene expression value was measured using the raw read count and the trimmed mean of M-values (TMM). Differential expression genes (DEGs) between different time points were identified by a generalized linear model (GLM) in R package edgeR (fold change>2 or <0.5, FDR<0.05). Significant temporal expression changes in times series data were identified by the regression strategy in R package maSigPro (v1.44.0).

Hierarchical clustering was used to cluster genes with similar expression patterns during *S. japonicum* infection. The number of clusters was manually selected to make the smallest cluster having more than 100 genes. Functional enrichment analysis of the interested gene sets was performed by DAVID, Gene Set Enrichment Analysis, and Ingenuity Pathway Analysis (IPA). The significant functional terms satisfied FDR<0.05. Ingenuity Upstream Regulator Analysis was used to identify the cascade of upstream transcriptional regulators that can explain the observed gene expression changes. To compare *M. fortis* and mouse, we only considered orthologous genes in these two species. The comparative transcriptome analysis was done at multiple levels: the number of DEGs, the Pearson correlation coefficients between gene expression fold change, enriched functions of DEGs, and the predicted upstream regulators.

## Discussion

*Microtus fortis* is not widely used in biomedical studies since it is distributed in particular regions. However, more and more research interests are raised to *M. fortis* due to its intrinsic resistance against *S. japonicum* infection and its potential as some disease models. To support further potential studies, we generate a draft reference genome for *M. fortis*. It will largely promote the further studies of gene functions and important traits in *M. fortis*. With decreased cost of third-generation sequencing, the reference genome can be improved to chromosome level by merging long-read sequencing and linkage mapping data.

The most attractive feature of *M. fortis* is the intrinsic resistance against schistosoma. There are several potential hypothesis: 1) *M. fortis* lack of genes that are necessary for the growth and development of schistosoma. 2) Compared to other species, *M. fortis* has a unique gene that prevent the development of schistosoma. 3) *M. fortis* has a special immune mechanism to prevent the development of schistosomulum.

After carefully investigating the sequence and evolution of *M. fortis* genes, we obtained 89 expanded and 102 contracted gene families. But the functional annotations of expanded genes and contracted genes seemed to have no direct relation with the growth and development of schistosoma. We found 532 positively selected genes (PSGs). Genes involving in innate immune progress were significantly enriched in PSGs. We also identified genes encoding immunoglobulin, T cell receptor and MHC. Their sequences will be extremely useful for further experimental studies, providing valuable resources for screening specific immune molecules against schistosoma.

More interestingly, comparative transcriptome analysis demonstrated that immune response was activated in *M. fortis* on the 10^th^ days post infection. Subsequent analysis of DEGs, immunoglobulins and upstream regulators discovered several possibly processes related to the protective immunity mechanism of *M. fortis* against schistosome (Figure 6). Leukocyte and other immune cells are recruited to the liver; Activated IRF7 initiates the induction of type I interferon, leads to the activation of JAK-STAT pathway and interferon stimulated genes; Antibodies (especially IgG3) are generated against schistosoma antigens; IgG binds to Fc-gamma receptor to induce phagocytosis. These different processes might work together to prevent the normal development of schistosoma. Previous studies showed that IgG antibodies of non-susceptible host rat could kill schistosomula of *S. mansoni* in *vitro* and in *vivo* [52], and IgG3 antibody of *M. fortis* could more effectively killed schistosomula than that of mouse [18]. Further investigation the functions of *M. fortis* IgG3 is very useful for discovering vaccines to protect people against schistosoma infection. Mouse has different immune responses after *S. japonicum* infection. Previous studies revealed that the dominant immunity of mouse is Th1 response at the 3^rd^~5^th^ weeks after *S. japonicum* infection, while Th2 response is generally to peak at the 8^th^ weeks [24]. Our results confirmed the activation of Th1 response at 28^th^~42^rd^ DPI in mouse, but Th2 response was not obvious due to the lack of data after 42^rd^ DPI. Th1 and Th2 activation pathways were also statistically significant for up-regulated genes at *M. fortis* 10^th^ DPI, but the expression of markers genes such as IFNG, TNF, IL4, IL10 and IL13 were not detected. Taken together, distinct mechanism of immune response is the most possible reason for the intrinsic resistance of *M. fortis* against schistosoma.

**Figure 6.**
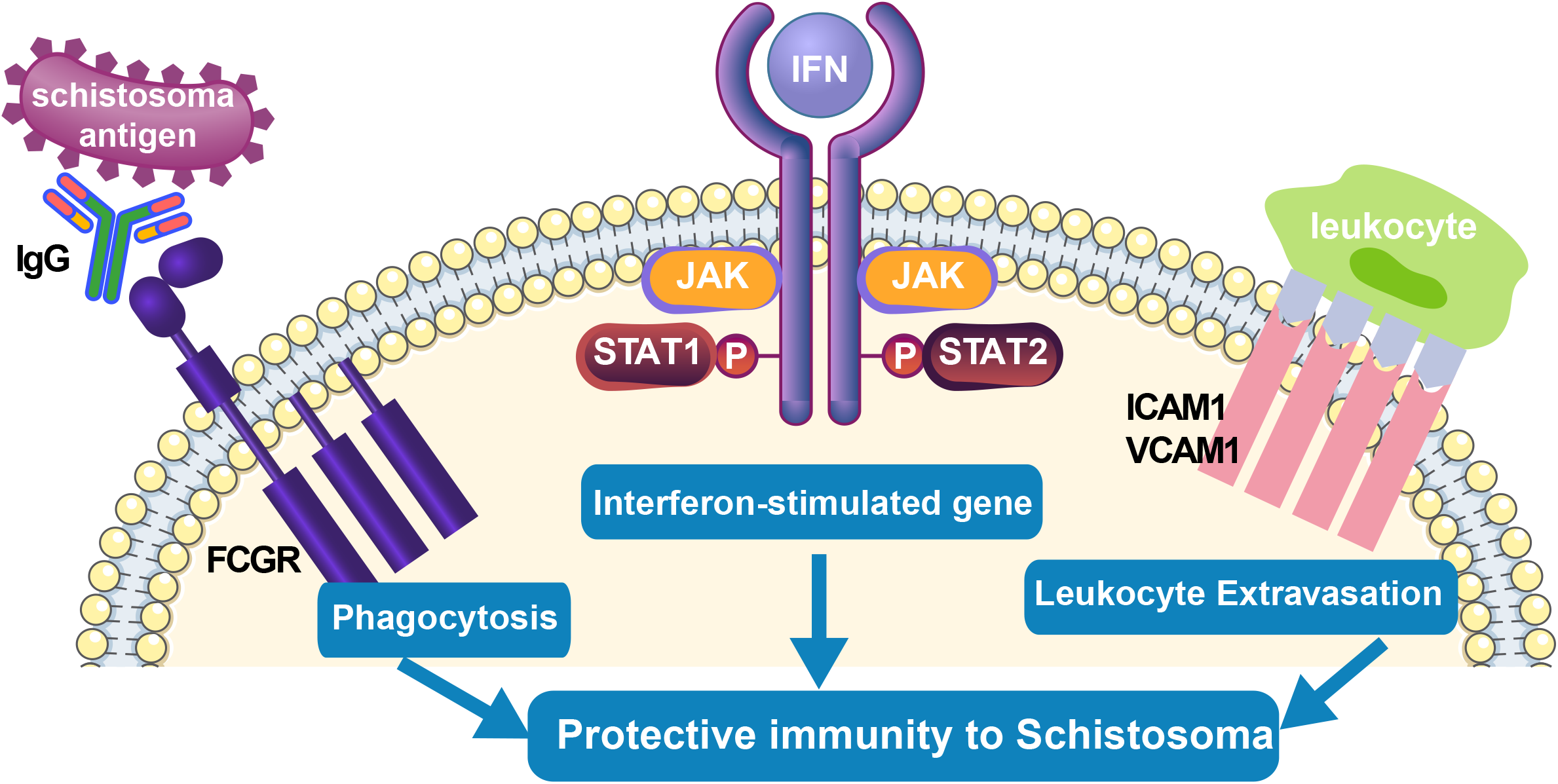
Potential immune processes related with the intrinsic resistance of *M. fortis*.

Intrinsic resistance of *M. fortis* against *S. japonicum* infection is a complex system. Our results comprehensively illustrated the dynamic expression patterns of different hosts after schistosoma infection, but the expression of some cytokines was not observed. One possible reason is that expression of these cytokines could not be detected in liver. Additionally, we proposed potential processes to explain the protective immunity mechanism. Our results provided new insights into the intrinsic resistance of *M. fortis* against schistosoma infection. However, further experimental studies are needed to validate the real contribution of these process, and there may be other biological processes involved. In future, we plan to do functional experiments to validate our hypothesis, we also plan to study the transcriptome of peripheral blood mononuclear cell, lymph gland and spleen to further investigate the immune response of *M. fortis*.

### Availability of Supporting Data and Materials

The *M. fortis* whole-genome project has been deposited at the DDBJ/ENA/GenBank under the accession NMRL00000000. Raw sequencing data has been submitted to the SRA (Sequence Read Archive) and NODE (National Omics Data Encyclopedia) databases. The accession numbers for DNA sequencing data are SRA:SRP111496 and NODE:OEP000443. Expression matrix of *M. fortis* and mouse have been deposited in the GEO database under accession GSE101654 and GSE101656.

**Table 1.** Global statistics of the *M. fortis* genome.

| Sequencing | Insert size | Total data (Gb) | Sequence coverage (x) |
| --- | --- | --- | --- |
| Libraries | 250, 500 bp | 167.37 | 70.0 |
|  | 2, 5, 10, 20 kb | 97.53 | 40.8 |
| Assembly | N50 (kb) | Number | Size (Gb) |
| Contigs | 60.7 | 9,910 | 2.1 |
| Scaffolds | 10,158 | 64 | 2.2 |
| Annotation | Number | Total length (Mb) | Percentage of genome (%) |
| GC content | / | 932.58 | 42.4 |
| Repeats | / | 666.41 | 30.2 |
| Genes | 21,867 | 459.89 | 20.9 |
| CDS | 21,867 | 30.96 | 1.4 |

## Authors’ Contributions

H.L., Z.Q.F, J.Y.X and Y.X.L designed the study. H.L. and Z.W. performed most of the computational analysis. J.Y.X., C.G., X.B. and J.F. kept a *M. fortis* colony. Z.Q.F, S.M.C. and J.J.L performed animal experiments. Q.Y.X. and B.P.M predict genes. Z.W. did evolution analysis. W.L.L aligned the genomes of multiple species. S.H did functional annotation. B.L. and W.H. provided valuable suggestions for the research plan and potential immune mechanism. Y.Y.L. and J.Y.L revised the manuscript.

## Acknowledgements

This work was supported by the Key Project in the National Science & Technology Pillar Program from the Ministry of Science and Technology (Grant No. 2015BAI09B04), the National Natural Science Foundation of China (31872256, 31472188), National Key Research and Development Program of China (2017YFD0501306), the Chinese Academy of Sciences (KFJ-STS-QYZD-126, ZDBS-SSW-DQC-02), CAS Youth Innovation Promotion Association, SA-SIBS Scholarship Program.

