## Supplementary for "Genome assembly and transcriptome analysis provide insights into the anti-schistosome mechanism of *Microtus fortis*"

Supplementary information

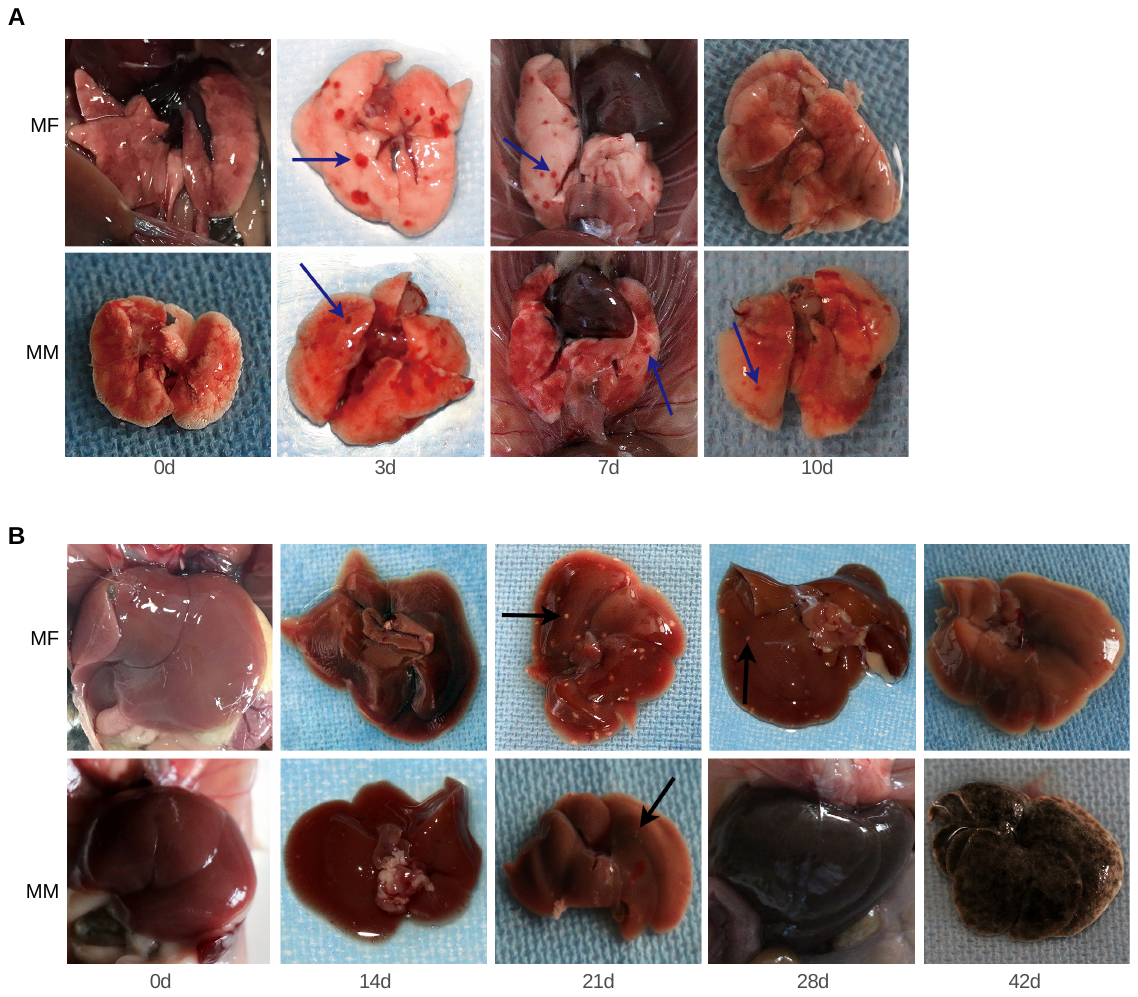

**Supplementary Figure 1**. Pictures of the lungs and livers from hosts infected with *S. japonicum*.

A) lungs. B) livers. The first line is the tissues prepared from S.*japonicum-*infected *M.fortis*. The second line is the tissues prepared from S.*japonicum-*infected *mice*. Each column represents a sampling point: pre-infection (0d) and post-infections (14 days, 21 days, 28 days and 42 days after infection). The arrows in Figure A show bleeding points on the lung surface, and the arrows in Figure B show white nodules on the liver surface.

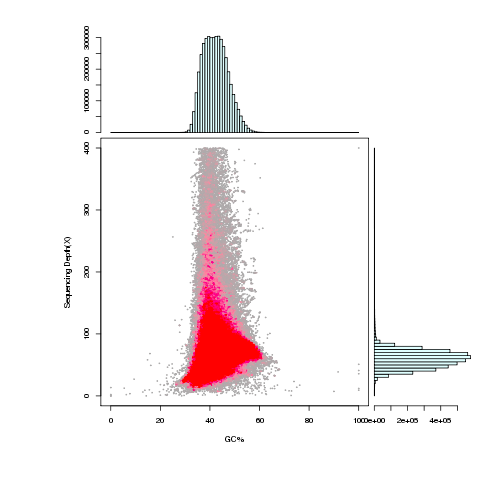

**Supplementary Figure 2**. GC content against the sequencing depth of *Microtus fortis* genome.

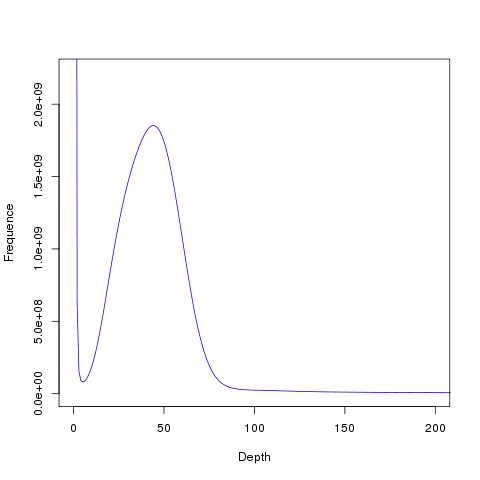

**Supplementary Figure 3.** Distribution of k-mer (k=39) sequences in the *Microtus fortis* genome.

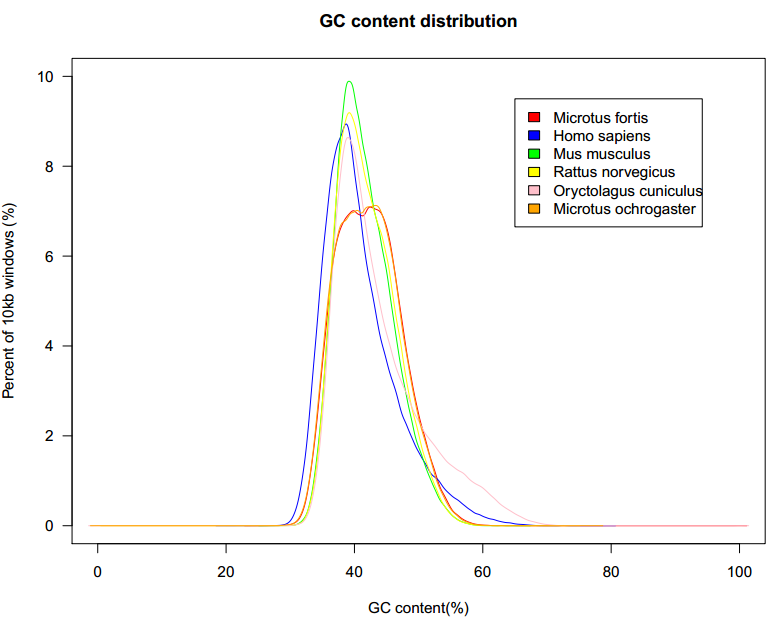

**Supplementary Figure 4.** The GC content for 10 kb, non-overlapping sliding windows across the *Microtus fortis* genome and other genomes.

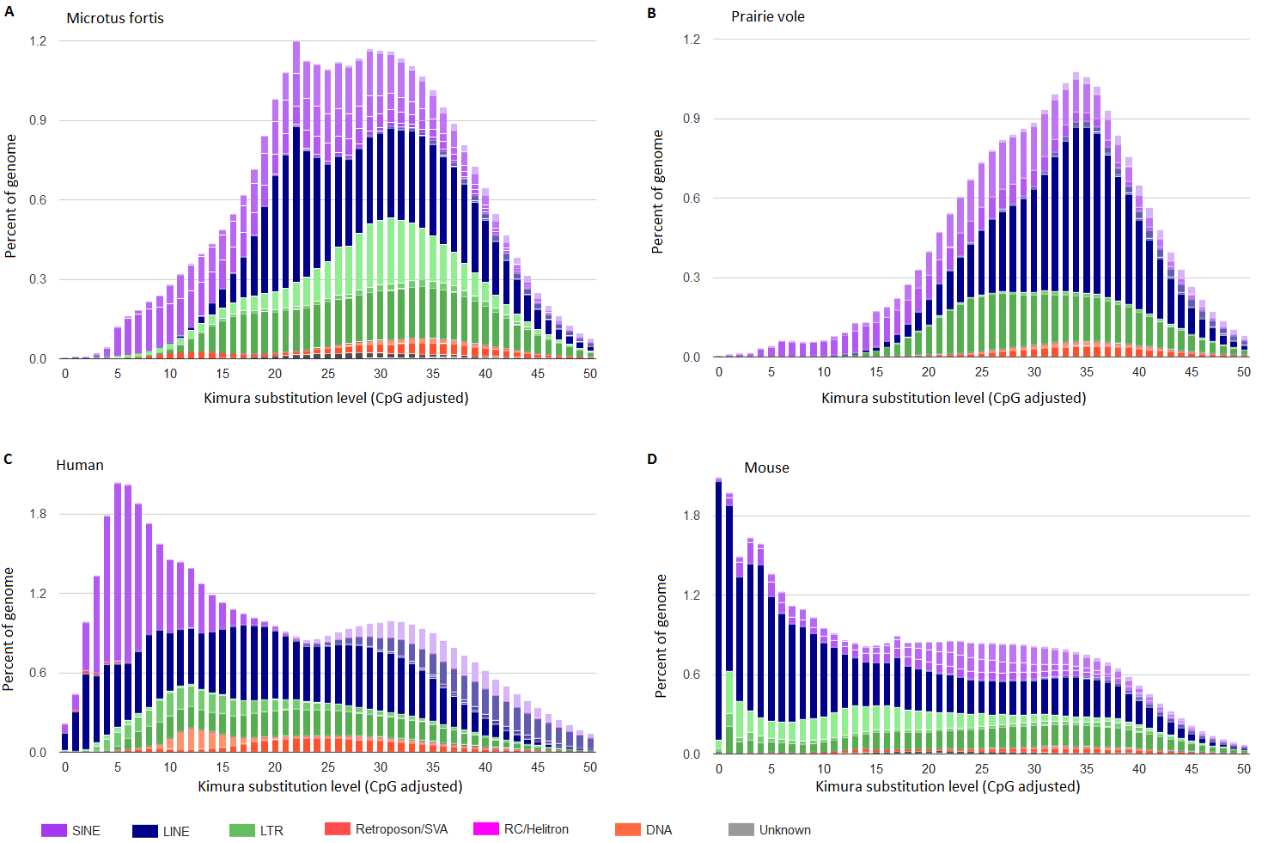

**Supplementary Figure 5.** The repeat landscapes.

It depicts the relative abundance of repeat classes in the genomes.

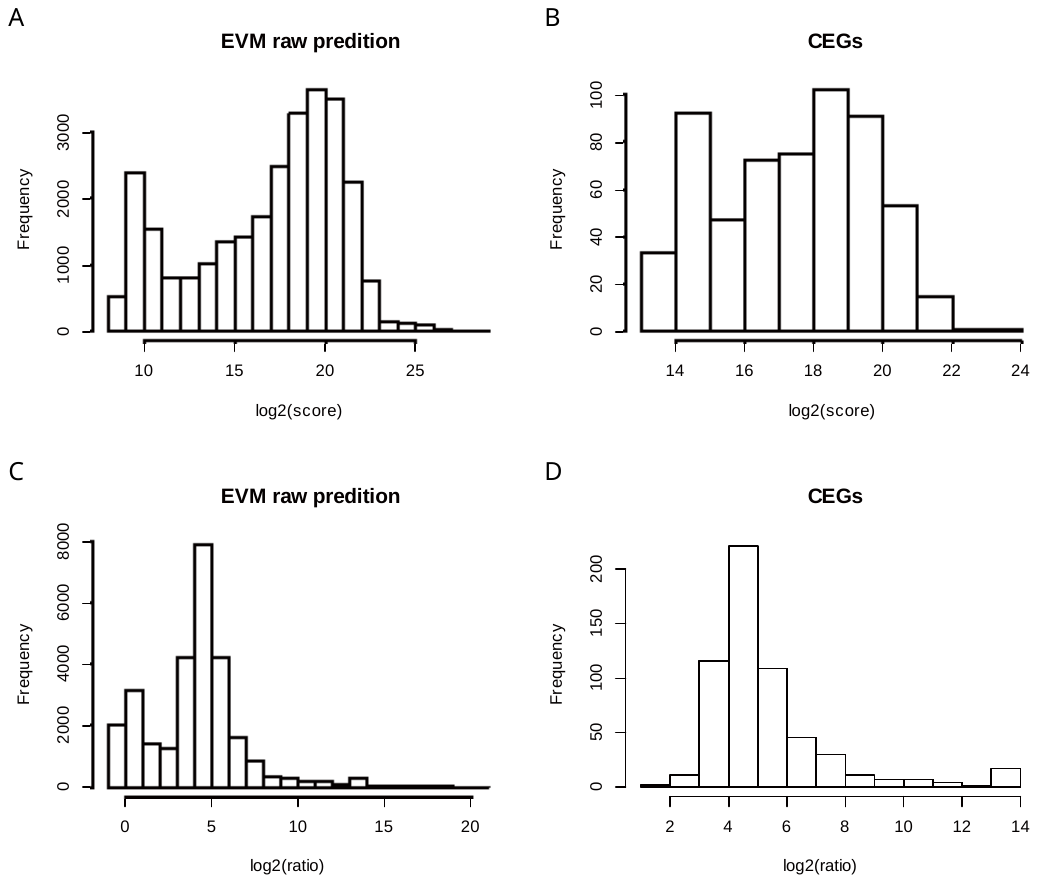

**Supplementary Figure 6.** Score of the raw EVM results.

EVidenceModeler (EVM) was used to combine *de novo* prediction, homology-based prediction, and RNA-Seq data into weighted consensus gene structures. It reports gene structures and scores the exon, intron, and intergenic region features. A), B) Distribution of log2(coding score) in EVM predictions and CEGs. C), D) Distribution of log2(coding/noncoding score ratio) in EVM predictions and CEGs. CEGs are core eukaryotic genes, which were regarded as positive control in the EVM results. EVM predictions with a coding score below 1024 and a coding/noncoding score ratio below 2 were eliminated.

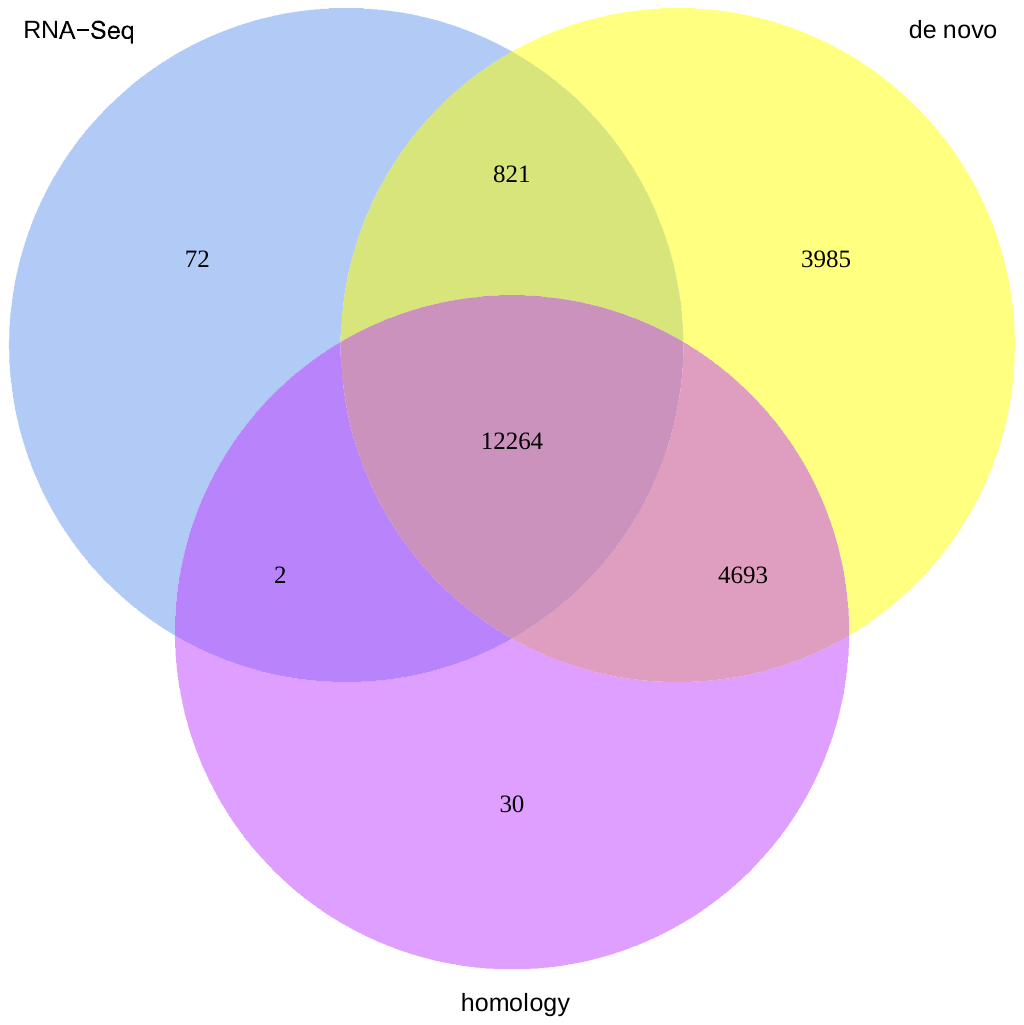

**Supplementary Figure 7.** Summary of evidence for the final gene models.

EVM was used to integrate *de novo* prediction, homology-based prediction and RNA-seq data. The low-quality gene models were filtered; the final gene models contain 21867 genes. Each gene model is supported by one or multiple evidence types.

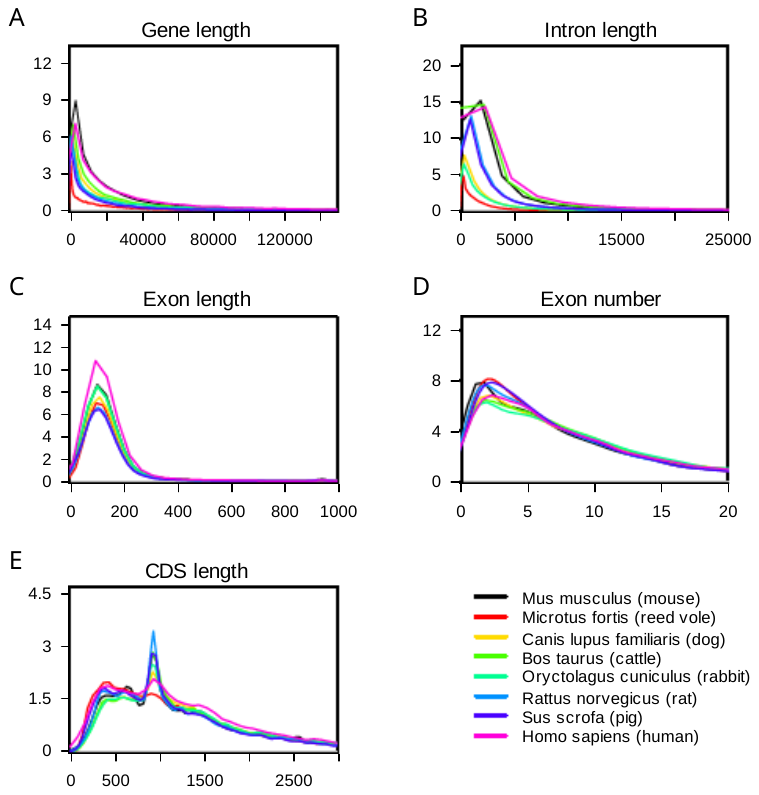

**Supplementary Figure 8.** Basic statistics of the predicted protein-coding genes.

*Microtus fortis* was compared with another seven species: human, dog, pig, cattle, mouse, rat and rabbit (from Ensembl release 75).

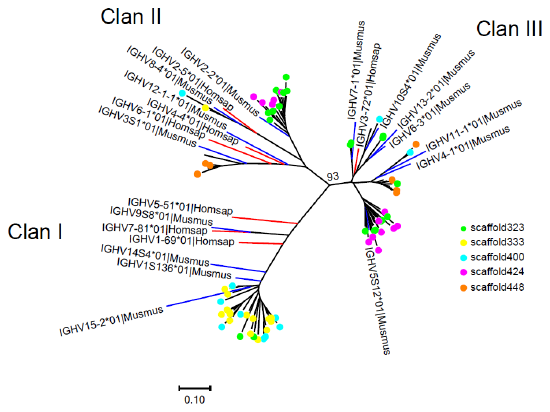

**Supplementary Figure 9**. Phylogenetic tree of *M. fortis* IGHV genes.

The human and mouse genes, each from one IGHV subgroup, are represented by red and blue branches, respectively. Similar to the human and mouse, three major clans are identified in *M. fortis*.

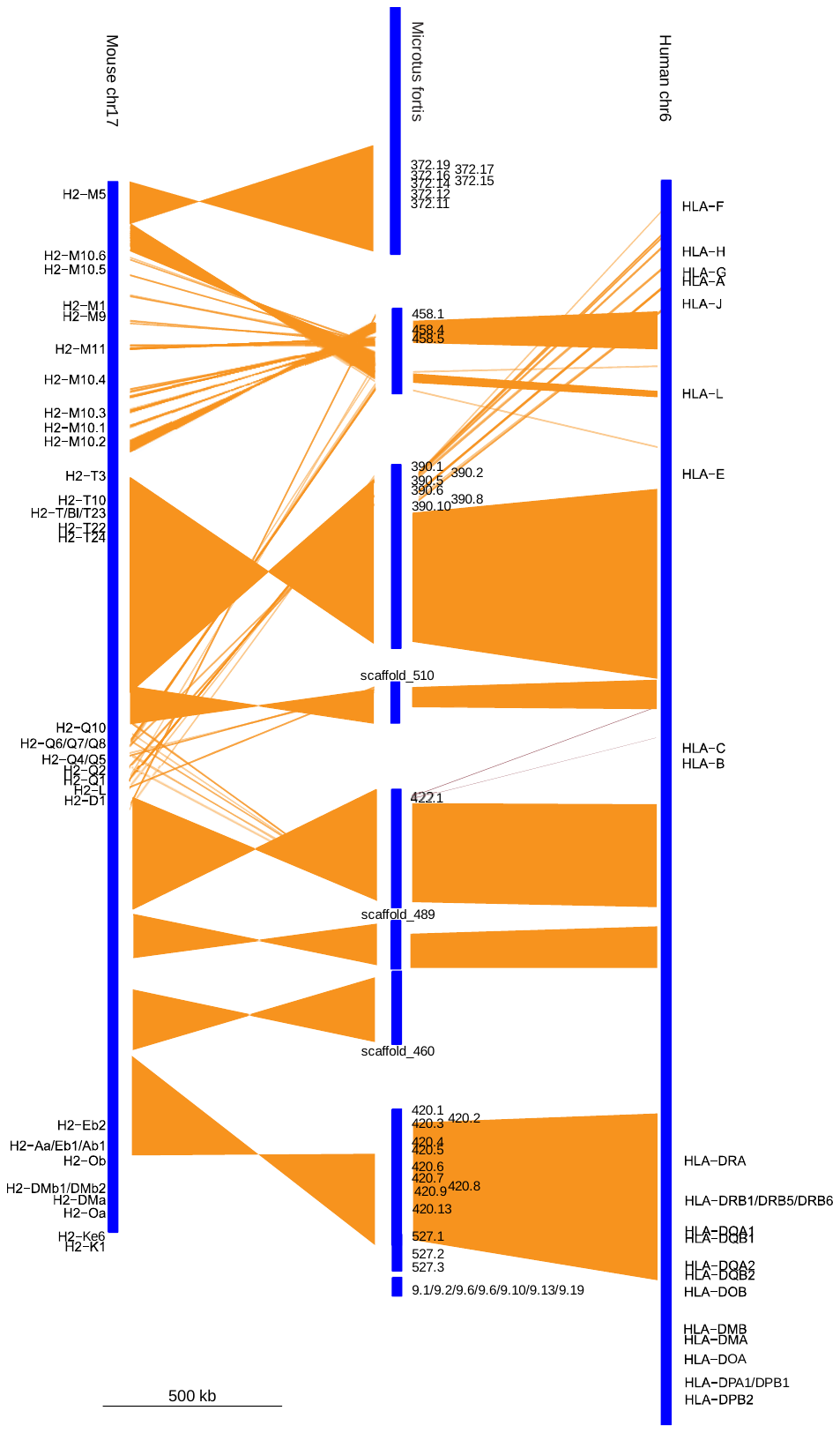

**Supplementary Figure 10.** Comparison of the structure and sequence of the MHC region in *M. fortis*, mouse and human.

Each genome is represented as a blue rectangle with tick marks showing the genomic position of MHC genes. Synteny regions between the two genomes are displayed as orange blocks. Annotation for human and mouse MHC region were retrieved from NCBI database. MHC genes of *M. fortis* are identified based on the annotation of KEGG, PFAM and GO databases.

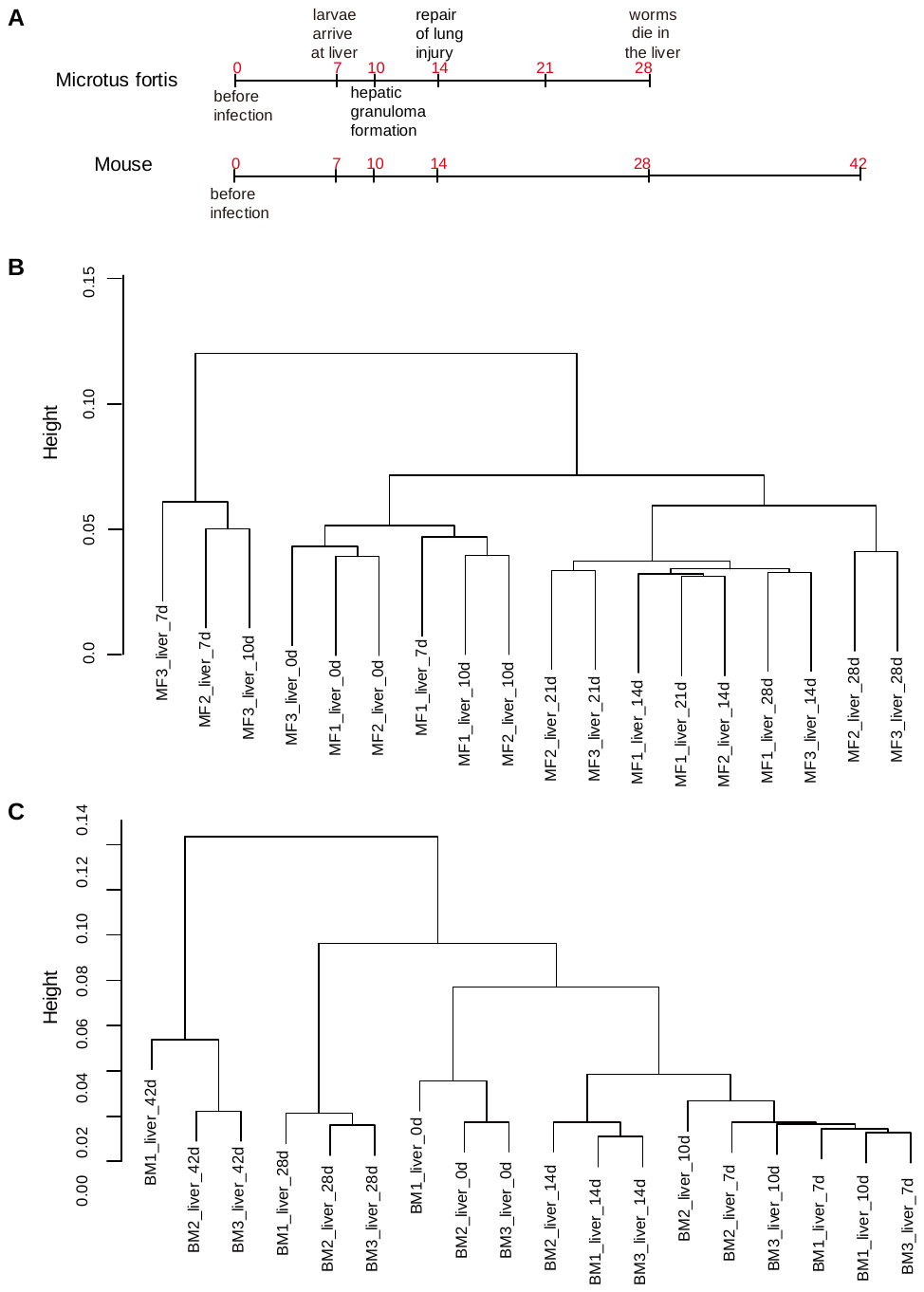

**Supplementary Figure 11.** Global expression patterns of *M. fortis* and mouse after infected with *Schistosoma japonicum*.

A) RNA-Seq experiment design. The start time “0” means before infection, other times are the days after infection. At each red labelled time points, three animals were subjected to RNA-Seq. B) Unsupervised clustering of *M. fortis* liver transcriptome. C) Unsupervised clustering of mouse liver transcriptome. All genes expressed in at least three of the samples with FPKM>=0.5 were used for the analysis. Similarity between paired samples was defined by spearman correlation coefficient.

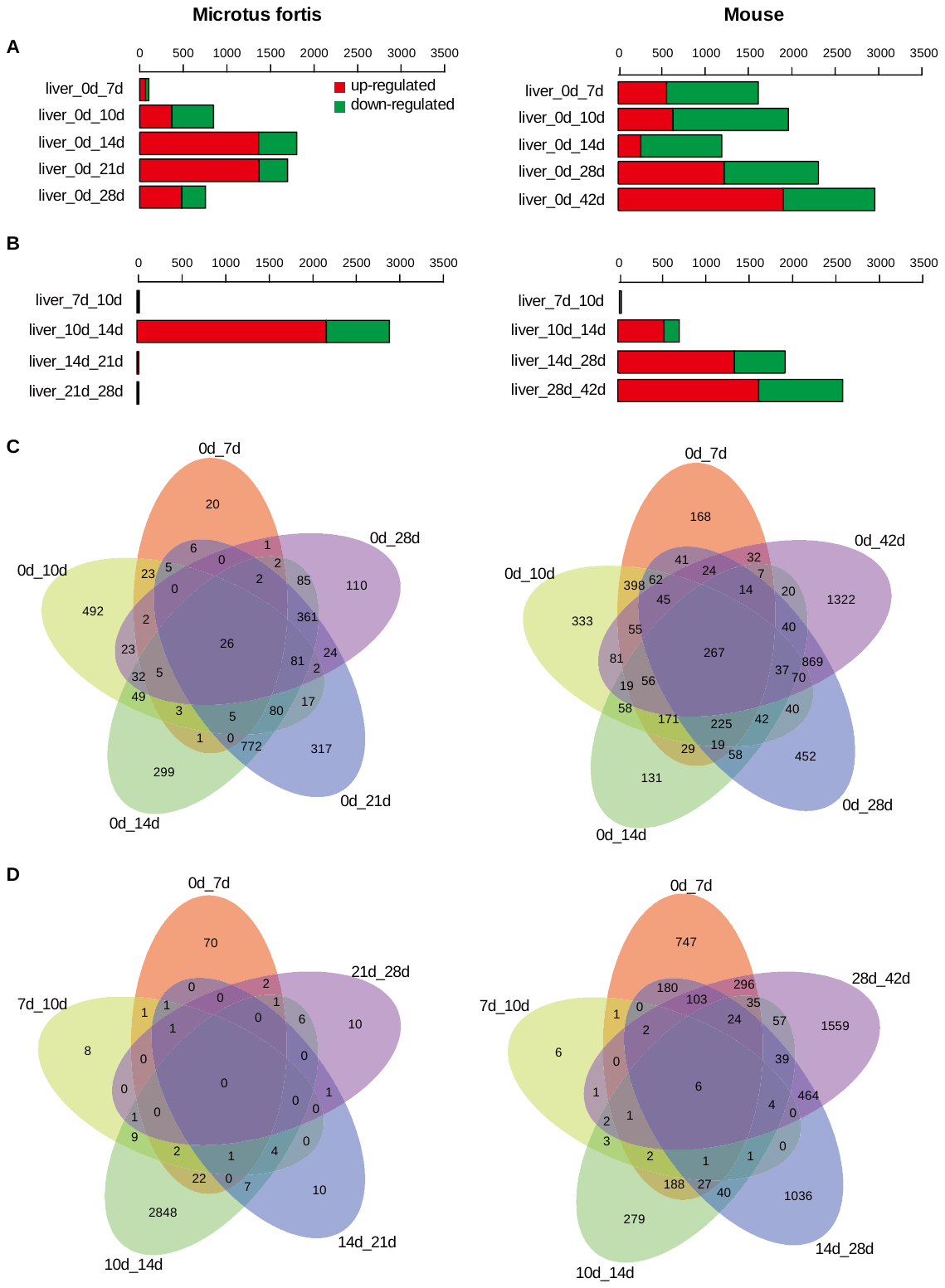

**Supplementary Figure 12. Overview of differential expression in** pairwise comparisons.

A) Number **of differential expression genes (DEGs) between pre-infection and post-infection. B) Number of DEGs between any two adjacent time points. C) Overlap between the numbers of DEGs between pre-infection and post-infection. D) Overlap between the numbers of DEGs any two adjacent time points. Differential expression analyses were performed by the generalized linear models (glms) in R package edgeR. Significantly differential genes were selected using false discovery rate (FDR) <0.05 and fold change (FC) >2 (or <0.5).**

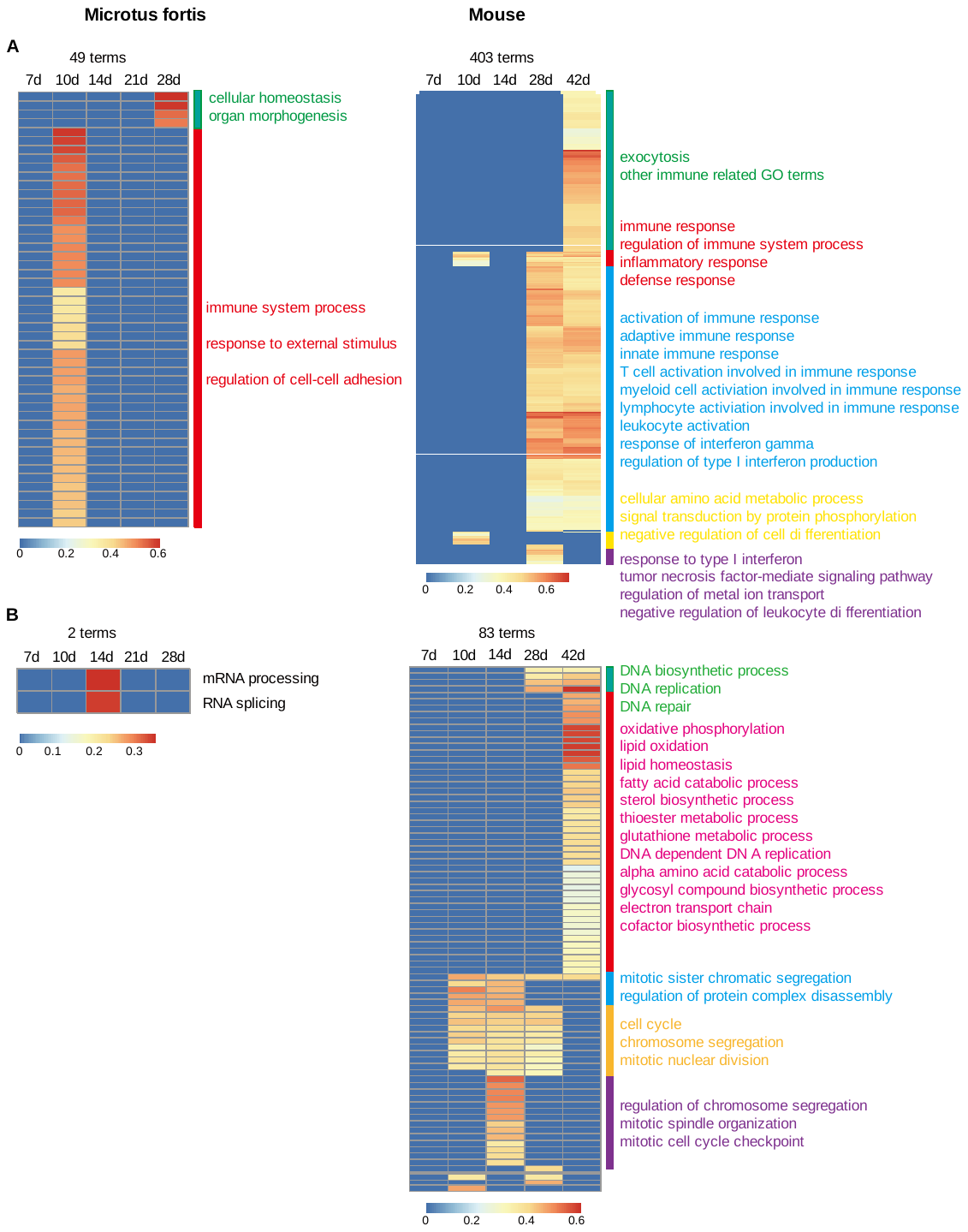

**Supplementary Figure 13.** Gene Ontology (GO) enrichment analysis of the differentially expressed genes (DEGs).

DEGs were significantly changed after Schistosome infection compared to pre-infection (FDR<0.05, FC>2 or <0.5). A) Up-regulated genes after infection. B) Down-regulated genes after infection. Enrichment analyses were performed by GSEA algorithm. GO biological process terms were obtained from MSigDB gene sets C5. GO terms with FDR<0.05 were regarded as significant, and only a part of representative terms are listed in the figure.

­

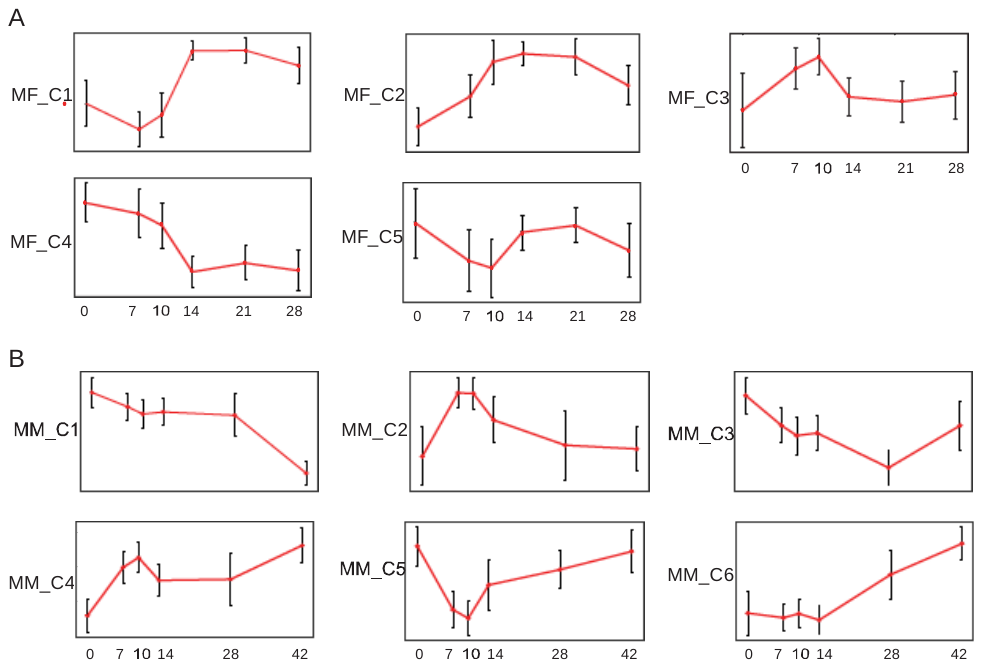

**Supplementary Figure 14.** Representative expression patterns of each DEG cluster.

A) *M. fortis*. B) Mouse.

**Supplementary Table 1**. Recovery rate of schistosoma adult worms in infected host.

| **Type of host** | **Host and reference** | **Recovery rate** |
| --- | --- | --- |
| Susceptible host | BALB/c mouse [1, 2] | 48%、60.6% |
| Susceptible host | C57BL mouse [3] | 57.7% ~ 84.8% |
| Susceptible host | *Mesocricetus auratus* (golden Hamster) [3] | 71.7% ~ 80.6% |
| Susceptible host | Jird [3] | 73.1% ~ 82.3% |
| Susceptible host | *Oryctolagus cuniculus* (rabbit) [3] | 69.3% ~ 80.8% |
| Susceptible host | *Canis familiaris* (dog) [4] | 59.0% |
| Susceptible host | *Macaca mulatta* (rhesue monkey) [5] | 35.9% ~ 78.3% |
| Susceptible host | *Bos Taurus* (yellow cattle) [6] | 54.0% ~ 57.0% |
| Susceptible host | *Capra hircus* (goat) [1] | 60.3% |
| Non-Susceptible host | *Bubalus bubalis* (water buffalo) [6] | 4.6% ~ 7.4% |
| Non-Susceptible host | Wistar rat [1] | 2% |
| Non-Susceptible host | *Swine* [4] | 8.5% |
| Non-permissive host | *Microtus fortis* (Reed vole) | 0% |

**Supplementary Table 2.** Genome sequencing strategy for the *M. fortis.*

| **Insert size** | **Original data** | | **Clean data after removing adapters and low-quality bases** | |
| --- | --- | --- | --- | --- |
|  | **Total bases (Gb)** | **Sequence depth (X)** | **Total bases (Gb)** | **Sequence depth (X)** |
| 250 bp | 83.81 | 35.07 | 83.11 | 34.77 |
| 500 bp | 83.56 | 34.96 | 83.33 | 34.87 |
| 2 kb | 30.78 | 12.88 | 30.34 | 12.69 |
| 5 kb | 33.17 | 13.88 | 31.74 | 13.28 |
| 10 kb | 24.18 | 10.12 | 22.01 | 9.21 |
| 20 kb | 9.4 | 3.93 | 9.11 | 3.81 |
| Total | 265 | 111 | 260 | 108 |

**Supplementary Table 3.** Estimation of the *M. fortis* genome size using K-mer analysis.

| **Kmer** | **K-mer number** | **K-mer depth** | **Genome** **size** (**Mbp**) | **Revised genome** **size (Mbp)** | **Heterozygous ratio (%)** | **Repeat** **(%)** |
| --- | --- | --- | --- | --- | --- | --- |
| 39 | 103,161,278,622 | 43 | 2,399 | 2,128 | 0.64 | 14.27 |

**Supplementary Table 4.** Summary of the *M. fortis* genome assembly.

|  | **length** | | **number** | |
| --- | --- | --- | --- | --- |
|  | **contig (bp)** | **scaffold (bp)** | **contig** | **scaffold** |
| Total | 2,104,733,352 | 2,201,563,765 | 78,011 | 10,542 |
| Longest | 560,751 | 48,647,192 | - | - |
| Number>=100bp | / | / | 78,011 | 10,542 |
| Number>=2kb | / | / | 65,298 | 5,162 |
| N50 | 60,779 | 10,158,217 | 9,910 | 64 |
| N60 | 47,875 | 8,235,182 | 13,810 | 88 |
| N70 | 36,579 | 5,548,218 | 18,837 | 121 |
| N80 | 25,602 | 3,808,250 | 25,694 | 169 |
| N90 | 14,864 | 1,779,544 | 36,282 | 251 |

**Supplementary Table 5.** Comparison of genome statistics across species.

|  | **Human** | **Mouse** | **Rat** | **Rabbit** | ***M. ochrogaster*** | ***M. fortis*** |
| --- | --- | --- | --- | --- | --- | --- |
| Genome size (Mb) | 3209.29 | 2730.87 | 2909.34 | 2737.33 | 2285.47 | 2201.56 |
| GC content (%) | 41.06 | 42.61 | 42.35 | 44.05 | 42.83 | 42.36 |

**Supplementary Table 6.** Coverage rate of core eukaryotic genes (CEGs) by CEGMA.

| **CEGMA**  (248 CEGs) | **Complete** | | **Partial** | |
| --- | --- | --- | --- | --- |
|  | Genes | Completeness (%) | Genes | Completeness (%) |
| In the assembled genomes | 230 | 92.74 | 238 | 95.97 |
| After gene annotation (21867 genes) | 236 | 95.16 | 245 | 98.79 |

**Supplementary Table 7.** Coverage rate of transcriptional fragments.

Transcriptional fragments were obtained from de novo RNA-Seq assembly by Trinity.

| **Dataset** | **Number** | **Total length (bp)** | **Sequences Covered by assembly (%)** | **with >90% sequence in one scaffold** | | **with >50% sequence in one scaffold** | |
| --- | --- | --- | --- | --- | --- | --- | --- |
|  |  |  |  | **Number** | **Percent (%)** | **Number** | **Percent (%)** |
| **>200bp** | 116,254 | 88,675,239 | 99.60 | 114,594 | 98.57 | 115,594 | 99.43 |
| **>500bp** | 40,110 | 66,356,406 | 99.68 | 39,893 | 99.46 | 39,975 | 99.66 |
| **>1k** | 21,958 | 53,810,187 | 99.79 | 21,897 | 99.72 | 21,910 | 99.78 |
| **>2k** | 10,999 | 38,110,062 | 99.88 | 10,984 | 99.86 | 10,986 | 99.88 |

**Supplementary Table 8.** General statistics of repeat elements.

| **Repeat type** | | **Proportion (%) in total nucleotide bases** | | | | | |
| --- | --- | --- | --- | --- | --- | --- | --- |
| **Class** | **Subclass** | **Human** | **Mouse** | **Rat** | **Rabbit** | **Prairie vole** | **Reed vole** |
| **SINE** |  | 12.79 | 7.46 | 6.62 | 19.14 | 6.00 | 9.10 |
|  | **5S-Deu-L2** | 0.01 | 0.00 | 0.00 | 0.01 | 0.01 | 0.00 |
|  | **Alu** | 10.03 | 2.43 | 1.56 | 0.00 | 2.79 | 3.05 |
|  | **B2** |  | 2.19 | 2.00 |  |  | 2.32 |
|  | **B4** |  | 2.14 | 2.01 |  | 2.53 | 2.70 |
|  | **ID** |  | 0.16 | 0.55 |  |  | 0.28 |
|  | **MIR** | 2.72 | 0.54 | 0.49 | 1.56 | 0.66 | 0.55 |
|  | **tRNA** | 0.01 | 0.00 | 0.01 | 0.01 | 0.01 | 0.00 |
|  | **tRNA-Deu** | 0.00 | 0.00 | 0.00 | 0.00 | 0.00 | 0.00 |
|  | **tRNA-RTE** | 0.02 | 0.00 | 0.00 | 0.01 | 0.00 | 0.00 |
|  | **tRNA-C** |  |  |  | 17.55 |  |  |
|  | **7SL** |  |  |  |  |  | 0.20 |
| **LINE** |  | 20.73 | 19.85 | 16.49 | 16.07 | 8.82 | 9.69 |
|  | **CR1** | 0.39 | 0.07 | 0.07 | 0.20 | 0.09 | 0.07 |
|  | **Dong-R4** | 0.00 | 0.00 | 0.00 | 0.00 | 0.00 | 0.00 |
|  | **Jockey** | 0.00 | 0.00 | 0.00 | 0.00 | 0.00 | 0.00 |
|  | **L1** | 16.62 | 19.34 | 16.03 | 14.11 | 8.19 | 9.19 |
|  | **L1-Tx1** | 0.00 | 0.00 | 0.00 | 0.00 | 0.00 | 0.00 |
|  | **L2** | 3.56 | 0.42 | 0.37 | 1.69 | 0.51 | 0.41 |
|  | **Penelope** | 0.00 | 0.00 | 0.00 | 0.00 | 0.00 | 0.00 |
|  | **RTE-BovB** | 0.04 | 0.01 | 0.01 | 0.02 | 0.02 | 0.01 |
|  | **RTE-X** | 0.12 | 0.01 | 0.01 | 0.05 | 0.01 | 0.01 |
| **LTR** |  | 8.84 | 11.70 | 9.11 | 4.23 | 4.60 | 9.79 |
|  | **ERV1** | 2.77 | 1.14 | 0.91 | 0.01 | 0.24 | 0.55 |
|  | **ERVK** | 0.30 | 4.81 | 3.36 | 0.80 |  | 3.92 |
|  | **ERVL** | 1.89 | 1.19 | 0.92 | 0.89 | 0.64 | 1.04 |
|  | **ERVL-MaLR** | 3.60 | 4.52 | 3.88 | 1.03 | 3.67 | 4.24 |
|  | **Gypsy** | 0.17 | 0.02 | 0.02 | 1.38 | 0.03 | 0.02 |
|  | **ERV-Lenti** |  |  |  | 0.06 |  |  |
|  | **Other** | 0.11 | 0.02 | 0.02 | 0.06 | 0.02 | 0.02 |
| **DNA** |  | 3.51 | 1.09 | 0.98 | 1.55 | 0.0091 | 1.29 |
|  | **Kolobok** | 0.00 | 0.00 | 0.00 | 0.00 | 0.00 | 0.00 |
|  | **MULE-MuDR** | 0.02 |  |  |  |  |  |
|  | **Merlin** | 0.00 |  |  |  |  |  |
|  | **PIF-Harbinger** | 0.00 | 0.00 | 0.00 | 0.00 | 0.00 | 0.00 |
|  | **PiggyBac** | 0.02 | 0.00 | 0.00 | 0.00 | 0.00 | 0.00 |
|  | **TcMar** | 0.01 | 0.00 | 0.00 | 0.01 | 0.00 | 0.00 |
|  | **TcMar-Mariner** | 0.09 | 0.01 | 0.01 | 0.01 | 0.01 | 0.01 |
|  | **TcMar-Pogo** | 0.00 | 0.00 | 0.00 | 0.00 | 0.00 | 0.00 |
|  | **TcMar-Tc1** | 0.00 | 0.00 | 0.00 | 0.00 | 0.00 | 0.00 |
|  | **TcMar-Tc2** | 0.05 | 0.01 | 0.01 | 0.03 | 0.01 | 0.02 |
|  | **TcMar-Tigger** | 1.22 | 0.20 | 0.18 | 0.36 | 0.23 | 0.25 |
|  | **hAT** | 0.05 | 0.02 | 0.01 | 0.03 | 0.02 | 0.02 |
|  | **hAT-Ac** | 0.02 | 0.02 | 0.01 | 0.02 | 0.02 | 0.01 |
|  | **hAT-Blackjack** | 0.11 | 0.03 | 0.03 | 0.06 | 0.03 | 0.04 |
|  | **hAT-Charlie** | 1.52 | 0.69 | 0.64 | 0.81 | 0.46 | 0.80 |
|  | **hAT-Tag1** | 0.02 | 0.01 | 0.00 | 0.01 | 0.01 | 0.01 |
|  | **hAT-Tip100** | 0.34 | 0.08 | 0.07 | 0.18 | 0.09 | 0.10 |
|  | **Harbinger** |  |  |  |  |  | 0.00 |
|  | **Other** | 0.04 | 0.02 | 0.02 | 0.03 | 0.03 | 0.03 |
| **RC** | **Helitron** | 0.01 | 0.00 | 0.00 | 0.01 | 0.00 | 0.00 |
| **Retroposon** | **SVA** | 0.14 |  |  |  |  |  |
| **Other** |  |  | 0.30 | 0.25 |  |  | 0.30 |
| **Unknown** |  | 0.03 | 0.10 | 1.15 | 0.02 | 0.02 | 0.09 |
| **Total** | | 46.07 | 40.53 | 34.65 | 41.03 | 20.36 | 30.27 |

**Supplementary Table 9.** Statistics of the predicted protein-coding genes.

| **Type** | **Tool** | **Number** | **Average transcript length (bp)** | **Average CDS length (bp)** | **Average exon per gene** | **Average exon length (bp)** | **Average intron length (bp)** |
| --- | --- | --- | --- | --- | --- | --- | --- |
| ***de novo* prediction** | AUGUSTUS | 24949 | 46558.51 | 1412.83 | 8.54 | 165.43 | 5987.22 |
|  | GENEID | 48395 | 26912.16 | 888.41 | 5.18 | 171.58 | 6218.14 |
|  | GlimmerHMM | 109960 | 10277.41 | 640.07 | 3.3 | 193.89 | 4187.91 |
|  | SNAP | 120303 | 10867.96 | 766.61 | 5.39 | 142.28 | 2301.96 |
|  | GeneMark-ES/ET | 50854 | 8929.54 | 870.82 | 5.87 | 148.31 | 1654.29 |
| **homology-based** | H_sapien | 19878 | 24381.05 | 1553.65 | 7.95 | 195.40 | 3283.90 |
|  | M_musculus | 22680 | 22566.75 | 1502.93 | 7.44 | 202.00 | 3270.63 |
|  | R_norvegicus | 23923 | 21523.72 | 1451.59 | 7.17 | 202.46 | 3253.29 |
|  | O_cuniculus | 20107 | 23648.93 | 1487.53 | 7.56 | 196.85 | 3379.93 |
|  | B_taurus | 21526 | 21943.35 | 1457.83 | 7.31 | 199.42 | 3246.35 |
|  | S_scrofa | 23374 | 19039.77 | 1323.98 | 6.73 | 196.80 | 3093.15 |
|  | C_familiaris | 19447 | 23987.16 | 1545.77 | 7.96 | 194.14 | 3223.29 |
| **supported by RNA-seq data** | NGSQC+tophat+cuffmerge | 19774 genes (36689 transcripts) | 35681.97 | - | 9.33 | 374.78 | 3828.18 |
| **Integration of coding genes** | EVidenceModeler (EVM) | 21867 | 21030.25 | 1407.77 | 8.01 | 175.73 | 2798.96 |

**Supplementary Table 10.** Statistics of protein-coding genes with annotation.

|  | **InterPro** | **KEGG** | | **Gene Ontology (GO)** | | |
| --- | --- | --- | --- | --- | --- | --- |
| **Number of genes** | **interproscan** | **gene** | **pathway** | **Molecular Function (MF)** | **Biological**  **Process (BP)** | **Cellular**  **Component (CC)** |
| 21867 | 21027 | 11523 | 6507 | 16135 | 16119 | 16914 |

**Supplementary Table 11**. Statistics of the predicted non-coding genes.

| **Coding genes** | **pseudogene** | **Non coding genes** | | | | |
| --- | --- | --- | --- | --- | --- | --- |
|  |  | **miRNA** | **tRNA** | **rRNA** | **lncRNA** | **snRNA** |
| 21867 | 1008 | 670 | 515 | 167 | 147 | 963 |

**Supplementary Table 12.** Statistics of homolog and ortholog families.

| **Species** | **Common name** | Taxonomy ID | **Gene number** | **Blast hits** | **Hmologous clusters** | **Orthologs** |
| --- | --- | --- | --- | --- | --- | --- |
| *Bubalus bubalis* | water buffalo | 89462 | 21532 | 3721821 | 16577 | 18484 |
| *Bos taurus* | cattle | 9913 | 21435 | 3921186 | 16523 | 18384 |
| *Canis lupus familiaris* | dog | 9615 | 19849 | 3541731 | 16271 | 17863 |
| *Cricetulus griseus* | Chinese hamster | 10029 | 21476 | 4110727 | 16375 | 18095 |
| *Cavia porcellus* | domestic guinea pig | 10141 | 20239 | 3676953 | 16088 | 17606 |
| *Dipodomys ordii* | Ord's kangaroo rat | 10020 | 19786 | 3480701 | 16044 | 17520 |
| *Heterocephalus glaber* | naked mole-rat | 10181 | 19798 | 3553229 | 16119 | 17647 |
| *Homo sapiens* | human | 9606 | 20366 | 3707064 | 16321 | 17768 |
| *Ictidomys tridecemlineatus* | thirteen-lined ground squirrel | 43179 | 19901 | 3757473 | 16245 | 17906 |
| *Mesocricetus auratus* | golden hamster | 10036 | 19987 | 3562981 | 16211 | 17794 |
| *Monodelphis domestica* | gray short-tailed opossum | 13616 | 20891 | 4355320 | 17279 | 16411 |
| *Microtus fortis* | reed vole | 100897 | 21867 | 3175037 | 15798 | 17310 |
| *Mus musculus* | house mouse | 10090 | 22459 | 4153061 | 16473 | 18085 |
| *Microtus ochrogaster* | prairie vole | 79684 | 20206 | 3679796 | 16320 | 17975 |
| *Nannospalax galili* | Upper Galilee mountains blind mole rat | 1026970 | 20882 | 3919160 | 16431 | 18041 |
| *Oryctolagus cuniculus* | rabbit | 9986 | 20597 | 3659706 | 15669 | 17162 |
| *Rattus norvegicus* | Norway rat | 10116 | 23584 | 4256178 | 16610 | 18290 |
| *Sus scrofa* | pig | 9823 | 24105 | 3996329 | 16184 | 18051 |

**Supplementary Table 13.** Innate immune genes (GO:0045087) that are under positive selection.

| **Symbol** | **Gene ID** | **P_value** |  | **Symbol** | **Gene ID** | **P_value** |
| --- | --- | --- | --- | --- | --- | --- |
| AXL | 394.8 | 4.68E-07 |  | IRF7 | 3.204 | 4.17E-02 |
| BPIFB1 | 27.329 | 2.16E-02 |  | ITGAM | 146.83 | 1.39E-02 |
| C7 | 37.17 | 4.10E-02 |  | LAG3 | 150.37 | 3.74E-03 |
| CD86 | 187.34 | 7.30E-04 |  | LCN2 | 72.277 | 4.01E-02 |
| CD96 | 4.256 | 1.53E-02 |  | MUC5AC | 3.234 | 1.96E-04 |
| CFB | 489.5 | 2.44E-03 |  | PGLYRP2 | 42.34 | 1.25E-02 |
| DRD2 | 89.119 | 5.61E-04 |  | PIK3CD | 88.159 | 2.55E-03 |
| ECSIT | 242.5 | 3.33E-02 |  | PTAFR | 26.290 | 2.89E-02 |
| EIF2AK2 | 136.40 | 1.07E-02 |  | REL | 7.417 | 1.37E-02 |
| ERAP1 | 163.50 | 6.26E-03 |  | RNASEL | 21.187 | 2.52E-03 |
| FGA | 102.50 | 2.94E-02 |  | SLAMF1 | 51.33 | 4.41E-03 |
| FGR | 26.279 | 5.33E-03 |  | STAT2 | 263.48 | 2.73E-02 |
| ICAM1 | 242.49 | 9.19E-03 |  | TLR9 | 36.118 | 2.60E-04 |
| IFNGR1 | 107.29 | 2.19E-02 |  | TMEM173 | 25.10 | 6.97E-03 |
| IL23R | 338.8 | 1.14E-02 |  | TNIP2 | 7.88 | 1.58E-03 |
| IRF3 | 161.73 | 1.96E-03 |  | TRIM26 | 458.9 | 1.53E-05 |
|  |  |  |  | TRIM27 | 5.317 | 1.36E-05 |

**Supplementary Table 14.** Annotation of immunoglobulin genes. Parentheses show sequences without stop codon.

| **Scaffold** | **Orient** | **Start** | **End** | **Locus** | **Gene type** | **Gene number** | **Pseudogenes** |
| --- | --- | --- | --- | --- | --- | --- | --- |
| scaffold_323 | - | 191640 | 925820 | IGH | V | 26 | 4 |
| scaffold_333 | + | 51388 | 915863 | IGH | V | 23 | 6 |
| scaffold_400 | + | 4530 | 350047 | IGH | V | 16 | 3 |
| scaffold_424 | - | 27195 | 321412 | IGH | V | 14 | 3 |
| scaffold_448 | - | 60975 | 298630 | IGH | V | 11 | 2 |
| scaffold_461 | + | 6134 | 225181 | IGH | V | 8 | 3 |
| scaffold_485 | - | 9847 | 136940 | IGH | V | 6 | 1 |
| scaffold_547 | - | 12217 | 75042 | IGH | V | 3 | 0 |
| scaffold_1655 | + | 3474 | 7151 | IGH | V | 2 | 2 |
| scaffold_182 | - | 3314007 | 3341778 | IGH | V | 2 | 1 |
| scaffold_2741 | + | 848 | 3192 | IGH | V | 2 | 1 |
| scaffold_1746 | + | 4636 | 4919 | IGH | V | 1 | 1 |
| scaffold_2094 | - | 7640 | 7939 | IGH | V | 1 | 0 |
| scaffold_221 | - | 75226 | 75520 | IGH | V | 1 | 1 |
| scaffold_2532 | + | 2232 | 2521 | IGH | V | 1 | 1 |
| scaffold_2557 | + | 2698 | 2994 | IGH | V | 1 | 0 |
| scaffold_6189 | + | 1440 | 1716 | IGH | V | 1 | 0 |
| scaffold_6996 | - | 835 | 1124 | IGH | V | 1 | 1 |
| scaffold_8948 | - | 410 | 710 | IGH | V | 1 | 0 |
| scaffold_988 | - | 347 | 647 | IGH | V | 1 | 0 |
| scaffold_452 | + | 34494 | 205564 | IGH | C | 6 | 0 |
|  |  |  |  |  | J | 7 | 0 |
| scaffold_407 | - | 5388 | 472092 | IGK | V | 11 | 1 |
| scaffold_409 | - | 7857 | 445170 | IGK | V | 11 | 3 |
| scaffold_464 | - | 35338 | 211557 | IGK | V | 10 | 2 |
| scaffold_440 | + | 41514 | 217507 | IGK | V | 9 | 0 |
| scaffold_457 | + | 31875 | 212163 | IGK | V | 8 | 4 |
| scaffold_611 | + | 12951 | 55022 | IGK | V | 7 | 4 |
| scaffold_436 | + | 3573 | 246020 | IGK | V | 6 | 1 |
| scaffold_473 | - | 40923 | 180729 | IGK | V | 6 | 1 |
| scaffold_436 | - | 146594 | 256691 | IGK | V | 5 | 0 |
| scaffold_440 | - | 735 | 190606 | IGK | V | 5 | 0 |
| scaffold_257 | + | 20741 | 150282 | IGK | V | 4 | 0 |
|  |  |  |  |  | C | 1 | 0 |
|  |  |  |  |  | J | 5 | 0 |
| scaffold_590 | - | 6757 | 53584 | IGK | V | 4 | 2 |
| scaffold_596 | - | 5510 | 47509 | IGK | V | 4 | 1 |
| scaffold_656 | + | 59 | 46686 | IGK | V | 4 | 3 |
| scaffold_338 | + | 727797 | 746462 | IGK | V | 3 | 1 |
| scaffold_569 | + | 24530 | 73311 | IGK | V | 3 | 0 |
| scaffold_587 | + | 34470 | 86895 | IGK | V | 3 | 2 |
| scaffold_708 | - | 10426 | 46589 | IGK | V | 3 | 2 |
| scaffold_789 | + | 268 | 18078 | IGK | V | 3 | 1 |
| scaffold_505 | - | 2678 | 49395 | IGK | V | 2 | 0 |
| scaffold_579 | + | 5139 | 65313 | IGK | V | 2 | 0 |
| scaffold_604 | + | 3285 | 5955 | IGK | V | 2 | 1 |
| scaffold_616 | - | 26260 | 57262 | IGK | V | 2 | 1 |
| scaffold_624 | - | 29790 | 45548 | IGK | V | 2 | 1 |
| scaffold_741 | - | 12383 | 26799 | IGK | V | 2 | 2 |
| scaffold_10 | - | 14476988 | 14477215 | IGK | V | 1 | 0 |
| scaffold_1130 | - | 1 | 233 | IGK | V | 1 | 0 |
| scaffold_1266 | - | 7476 | 7767 | IGK | V | 1 | 1 |
| scaffold_1542 | + | 1440 | 1747 | IGK | V | 1 | 1 |
| scaffold_1946 | + | 209 | 510 | IGK | V | 1 | 0 |
| scaffold_2015 | - | 3507 | 3794 | IGK | V | 1 | 1 |
| scaffold_2052 | - | 547 | 853 | IGK | V | 1 | 0 |
| scaffold_2147 | - | 6496 | 6796 | IGK | V | 1 | 0 |
| scaffold_2223 | + | 267 | 574 | IGK | V | 1 | 0 |
| scaffold_25 | - | 4069832 | 4070073 | IGK | V | 1 | 1 |
| scaffold_2588 | + | 12358 | 12618 | IGK | V | 1 | 0 |
| scaffold_3623 | - | 1267 | 1553 | IGK | V | 1 | 1 |
| scaffold_3714 | - | 420 | 726 | IGK | V | 1 | 0 |
| scaffold_3770 | - | 270 | 628 | IGK | V | 1 | 0 |
| scaffold_4383 | - | 316 | 619 | IGK | V | 1 | 0 |
| scaffold_464 | + | 216473 | 216777 | IGK | V | 1 | 0 |
| scaffold_5745 | + | 217 | 462 | IGK | V | 1 | 1 |
| scaffold_61 | + | 5991225 | 5991525 | IGK | V | 1 | 1 |
| scaffold_646 | + | 9425 | 9728 | IGK | V | 1 | 0 |
| scaffold_782 | + | 6618 | 6844 | IGK | V | 1 | 1 |
| scaffold_813 | - | 1 | 258 | IGK | V | 1 | 0 |
| scaffold_816 | + | 9803 | 10104 | IGK | V | 1 | 0 |
| scaffold_906 | + | 18025 | 18329 | IGK | V | 1 | 0 |
| scaffold_9266 | + | 317 | 605 | IGK | V | 1 | 0 |
| scaffold_966 | + | 158 | 446 | IGK | V | 1 | 1 |
| scaffold_445 | - | 33351 | 205714 | IGL | V | 13 | 3 |
| scaffold_334 | - | 51004 | 514278 | IGL | V | 9 | 4 |
|  |  |  |  |  | C | 2 | 0 |
|  |  |  |  |  | J | 2 | 0 |
| scaffold_672 | - | 4876 | 25485 | IGL | V | 3 | 1 |
| scaffold_465 | + | 110175 | 135190 | IGL | V | 2 | 0 |
| scaffold_161 | + | 3910780 | 3911064 | IGL | V | 1 | 1 |
| scaffold_219 | - | 1293922 | 1294227 | IGL | V | 1 | 0 |
| scaffold_239 | - | 189594 | 189821 | IGL | V | 1 | 1 |
| scaffold_271 | - | 935 | 1234 | IGL | V | 1 | 1 |
| scaffold_36 | - | 3133362 | 3133636 | IGL | V | 1 | 1 |
| scaffold_56 | - | 45118 | 45411 | IGL | V | 1 | 1 |

**Supplementary Table 15.** Annotation of T-cell receptor genes. Parentheses show sequences without stop codon.

| **Scaffold** | **Orient** | **Start** | **End** | **Type** | **Gene type** | **Gene number** | **Pseudogenes** |
| --- | --- | --- | --- | --- | --- | --- | --- |
| scaffold_87 | + | 1009 | 1056843 | TRA/TRD | V | 41 | 4 |
|  |  |  |  |  | C | 2 | 0 |
|  |  |  |  |  | J | 47 | 0 |
| scaffold_347 | - | 7370 | 537797 | TRA/TRD | V | 40 | 2 |
|  |  |  |  |  | C | 2 | 0 |
|  |  |  |  |  | J | 45 | 0 |
| scaffold_2426 | + | 212 | 2212 | TRA | V | 2 | 1 |
| scaffold_258 | - | 43007 | 59856 | TRA | V | 2 | 1 |
| scaffold_534 | - | 82994 | 85003 | TRA | V | 2 | 1 |
| scaffold_2659 | - | 1308 | 1581 | TRA | V | 1 | 1 |
| scaffold_2844 | - | 2851 | 3130 | TRA | V | 1 | 0 |
| scaffold_3906 | + | 539 | 812 | TRA | V | 1 | 1 |
| scaffold_9245 | + | 207 | 466 | TRA | V | 1 | 1 |
| scaffold_57 | + | 9706856 | 10273072 | TRB | V | 34 | 4 |
|  |  |  |  |  | C | 2 | 0 |
|  |  |  |  |  | J | 12 | 0 |
| scaffold_1763 | - | 5828 | 6114 | TRD | V: | 1 | 0 |
| scaffold_5 | - | 12062957 | 12120167 | TRG | V | 6 | 1 |
|  |  |  |  |  | C | 1 | 0 |
|  |  |  |  |  | J | 2 | 0 |

**Supplementary Table 16.** List of annotated MHC genes of *M. fortis.*

| **Gene ID** | **Scaffold** | **Start** | **End** | **Domain** |
| --- | --- | --- | --- | --- |
| 372.11 | scaffold_372 | 152724 | 155831 | PF00129 (Class I Histocompatibility antigen, domains alpha 1 and 2) |
| 372.12 | scaffold_372 | 167750 | 174637 | PF00129 (Class I Histocompatibility antigen, domains alpha 1 and 2) |
| 372.14 | scaffold_372 | 203527 | 212070 | PF00129 (Class I Histocompatibility antigen, domains alpha 1 and 2) |
| 372.15 | scaffold_372 | 213591 | 221372 | PF00129 (Class I Histocompatibility antigen, domains alpha 1 and 2) |
| 372.16 | scaffold_372 | 222616 | 226660 | PF00129 (Class I Histocompatibility antigen, domains alpha 1 and 2) |
| 372.17 | scaffold_372 | 232318 | 233889 | PF00129 (Class I Histocompatibility antigen, domains alpha 1 and 2) |
| 372.19 | scaffold_372 | 246093 | 263097 | PF00129 (Class I Histocompatibility antigen, domains alpha 1 and 2) |
| 390.1 | scaffold_390 | 4576 | 7568 | PF00129 (Class I Histocompatibility antigen, domains alpha 1 and 2) |
| 390.10 | scaffold_390 | 118443 | 119519 | PF00129 (Class I Histocompatibility antigen, domains alpha 1 and 2) |
| 390.2 | scaffold_390 | 19293 | 26214 | PF00129 (Class I Histocompatibility antigen, domains alpha 1 and 2) |
| 390.5 | scaffold_390 | 45635 | 48910 | PF00129 (Class I Histocompatibility antigen, domains alpha 1 and 2) |
| 390.6 | scaffold_390 | 70359 | 78796 | PF00129 (Class I Histocompatibility antigen, domains alpha 1 and 2) |
| 390.8 | scaffold_390 | 94927 | 101480 | PF00129 (Class I Histocompatibility antigen, domains alpha 1 and 2) |
| 420.1 | scaffold_420 | 265 | 8212 | PF00969 (Class II histocompatibility antigen, beta domain) |
| 420.13 | scaffold_420 | 279594 | 292296 | PF00969 (Class II histocompatibility antigen, beta domain) |
| 420.2 | scaffold_420 | 21607 | 30109 | PF00969 (Class II histocompatibility antigen, beta domain) |
| 420.3 | scaffold_420 | 40272 | 44367 | PF00993 (Class II histocompatibility antigen, alpha domain) |
| 420.4 | scaffold_420 | 91862 | 97885 | PF00969 (Class II histocompatibility antigen, beta domain) |
| 420.5 | scaffold_420 | 116031 | 120528 | PF00993 (Class II histocompatibility antigen, alpha domain) |
| 420.6 | scaffold_420 | 161792 | 167736 | PF00969 (Class II histocompatibility antigen, beta domain) |
| 420.7 | scaffold_420 | 187056 | 205394 | PF00969 (Class II histocompatibility antigen, beta domain) |
| 420.8 | scaffold_420 | 212162 | 226173 | PF00969 (Class II histocompatibility antigen, beta domain) |
| 420.9 | scaffold_420 | 230618 | 234562 | PF00993 (Class II histocompatibility antigen, alpha domain) |
| 422.1 | scaffold_422 | 21603 | 30203 | PF00129 (Class I Histocompatibility antigen, domains alpha 1 and 2) |
| 458.1 | scaffold_458 | 14458 | 18160 | PF00129 (Class I Histocompatibility antigen, domains alpha 1 and 2) |
| 458.4 | scaffold_458 | 67005 | 70384 | PF00129 (Class I Histocompatibility antigen, domains alpha 1 and 2) |
| 458.5 | scaffold_458 | 85917 | 89021 | PF00129 (Class I Histocompatibility antigen, domains alpha 1 and 2) |
| 522.3 | scaffold_522 | 65347 | 69807 | PF00129 (Class I Histocompatibility antigen, domains alpha 1 and 2) |
| 522.4 | scaffold_522 | 74152 | 88272 | PF00129 (Class I Histocompatibility antigen, domains alpha 1 and 2) |
| 522.5 | scaffold_522 | 95542 | 97501 | PF00129 (Class I Histocompatibility antigen, domains alpha 1 and 2) |
| 527.1 | scaffold_527 | 3366 | 8096 | PF00993 (Class II histocompatibility antigen, alpha domain) |
| 527.2 | scaffold_527 | 54552 | 59705 | PF00969 (Class II histocompatibility antigen, beta domain) |
| 527.3 | scaffold_527 | 89865 | 96937 | PF00993 (Class II histocompatibility antigen, alpha domain) |
| 544.1 | scaffold_544 | 443 | 4004 | PF00129 (Class I Histocompatibility antigen, domains alpha 1 and 2) |
| 544.3 | scaffold_544 | 32864 | 43441 | PF00129 (Class I Histocompatibility antigen, domains alpha 1 and 2) |
| 544.4 | scaffold_544 | 76012 | 92025 | PF00129 (Class I Histocompatibility antigen, domains alpha 1 and 2) |
| 675.2 | scaffold_675 | 30006 | 38948 | PF00129 (Class I Histocompatibility antigen, domains alpha 1 and 2) |
| 679.1 | scaffold_679 | 19788 | 31368 | PF00129 (Class I Histocompatibility antigen, domains alpha 1 and 2) |
| 679.2 | scaffold_679 | 40708 | 44522 | PF00129 (Class I Histocompatibility antigen, domains alpha 1 and 2) |
| 9.1 | scaffold_9 | 11989 | 15814 | PF00969 (Class II histocompatibility antigen, beta domain) |
| 9.10 | scaffold_9 | 193934 | 196463 | PF00993 (Class II histocompatibility antigen, alpha domain) |
| 9.11 | scaffold_9 | 203122 | 206277 | PF00993 (Class II histocompatibility antigen, alpha domain) |
| 9.13 | scaffold_9 | 214440 | 218338 | PF00969 (Class II histocompatibility antigen, beta domain) |
| 9.19 | scaffold_9 | 281805 | 284833 | PF00129 (Class I Histocompatibility antigen, domains alpha 1 and 2) |
| 9.2 | scaffold_9 | 55511 | 68147 | PF00969 (Class II histocompatibility antigen, beta domain) |
| 9.6 | scaffold_9 | 139251 | 145909 | PF00969 (Class II histocompatibility antigen, beta domain) |
| 9.7 | scaffold_9 | 153511 | 156803 | PF00993 (Class II histocompatibility antigen, alpha domain) |

**Supplementary Table 17.** Summary of RNA-Seq mapping and identified genes.

|  | **Raw paired reads** | **High quality paired reads** | **percentage of high-quality reads (%)** | **number of genes (FPKM>1)** |
| --- | --- | --- | --- | --- |
| **Mixed_tissues** | 44474320 | 41803442 | 93.99 | 15252 |
| **MF1_liver_0d** | 21986557 | 20195855 | 91.86 | 11354 |
| **MF1_liver_10d** | 20594250 | 19194173 | 93.20 | 11629 |
| **MF1_liver_14d** | 27843583 | 27791136 | 99.81 | 12299 |
| **MF1_liver_21d** | 21505583 | 21480374 | 99.88 | 12517 |
| **MF1_liver_28d** | 19666022 | 19643921 | 99.89 | 12589 |
| **MF1_liver_7d** | 24484333 | 22687328 | 92.66 | 11807 |
| **MF2_liver_10d** | 25375481 | 23566432 | 92.87 | 11123 |
| **MF2_liver_14d** | 27988785 | 27934452 | 99.81 | 12350 |
| **MF2_liver_21d** | 19951198 | 19925637 | 99.87 | 12392 |
| **MF2_liver_28d** | 19982263 | 18580215 | 92.98 | 11633 |
| **MF2_liver_7d** | 21180103 | 19671238 | 92.88 | 10742 |
| **MF3_liver_10d** | 20003647 | 18708566 | 93.53 | 11110 |
| **MF3_liver_14d** | 21712503 | 21687854 | 99.89 | 12565 |
| **MF3_liver_21d** | 19144353 | 19122128 | 99.88 | 12533 |
| **MF3_liver_28d** | 21466780 | 19935309 | 92.87 | 11607 |
| **MF3_liver_7d** | 22741251 | 21292693 | 93.63 | 10008 |
| **MM1_liver_0d** | 21153991 | 19214818 | 90.83 | 10566 |
| **MM1_liver_10d** | 39707685 | 37978600 | 95.65 | 9936 |
| **MM1_liver_14d** | 21429012 | 19959558 | 93.14 | 10155 |
| **MM1_liver_28d** | 21377860 | 18945000 | 88.62 | 10299 |
| **MM1_liver_42d** | 34515696 | 34515607 | 100.00 | 11430 |
| **MM1_liver_7d** | 30023810 | 28668359 | 95.49 | 9801 |
| **MM2_liver_0d** | 25049429 | 22796560 | 91.01 | 10805 |
| **MM2_liver_10d** | 29305417 | 27595930 | 94.17 | 9977 |
| **MM2_liver_14d** | 19969068 | 18606839 | 93.18 | 10305 |
| **MM2_liver_28d** | 21846578 | 20326513 | 93.04 | 10498 |
| **MM2_liver_42d** | 24582802 | 24582588 | 100.00 | 11310 |
| **MM2_liver_7d** | 26193145 | 25080919 | 95.75 | 10048 |
| **MM3_liver_0d** | 22144883 | 20062110 | 90.59 | 10767 |
| **MM3_liver_10d** | 31583396 | 29627510 | 93.81 | 9826 |
| **MM3_liver_14d** | 20101042 | 18668106 | 92.87 | 10186 |
| **MM3_liver_28d** | 21153986 | 19694232 | 93.10 | 10715 |
| **MM3_liver_42d** | 33481399 | 33480960 | 100.00 | 11546 |
| **MM3_liver_7d** | 32018860 | 30666231 | 95.78 | 9896 |

**Supplementary Table 18.** Comparison of the DEGs at the same time points between *M. fortis* and mouse.

This analysis only used genes with orthologs in *M. fortis* and mouse. Significantly genes were selected by FDR<=0.05. MF: *M. fortis*.

| 0d_7d | | mouse | | |  | 0d_10d | | mouse | | |
| --- | --- | --- | --- | --- | --- | --- | --- | --- | --- | --- |
|  |  | Up | Down | Unchanged |  |  |  | Up | Down | Unchanged |
| MF | Up | 7 | 4 | 31 |  | MF | Up | 47 | 51 | 395 |
|  | Down | 1 | 5 | 17 |  |  | Down | 167 | 100 | 458 |
|  | Unchanged | 1635 | 1143 | 8696 |  |  | Unchanged | 1316 | 1172 | 7840 |
| 0d_14d | | mouse | | |  | 0d_28d | | mouse | | |
|  |  | Up | Down | Unchanged |  |  |  | Up | Down | Unchanged |
| MF | Up | 133 | 111 | 1512 |  | MF | Up | 61 | 123 | 485 |
|  | Down | 31 | 127 | 1087 |  |  | Down | 46 | 67 | 400 |
|  | Unchanged | 414 | 581 | 7572 |  |  | Unchanged | 1195 | 1080 | 8111 |
